# Reconstitution of eukaryotic chromosomes and manipulation of DNA N6-methyladenine alters chromatin and gene expression

**DOI:** 10.1101/475384

**Authors:** Leslie Y. Beh, Galia T. Debelouchina, Derek M. Clay, Robert E. Thompson, Kelsi A. Lindblad, Elizabeth R. Hutton, John R. Bracht, Robert P. Sebra, Tom W. Muir, Laura F. Landweber

## Abstract

DNA N6-adenine methylation (6mA) has recently been reported in diverse eukaryotes, spanning unicellular organisms to metazoans. Yet the functional significance of 6mA remains elusive due to its low abundance, difficulty of manipulation within native DNA, and lack of understanding of eukaryotic 6mA writers. Here, we report a novel DNA 6mA methyltransferase in ciliates, termed MTA1. The enzyme contains an MT-A70 domain but is phylogenetically distinct from all known RNA and DNA methyltransferases. Disruption of MTA1 *in vivo* leads to the genome-wide loss of 6mA in asexually growing cells and abolishment of the consensus ApT dimethylated motif. Genes exhibit subtle changes in chromatin organization or RNA expression upon loss of 6mA, depending on their starting methylation level. Mutants fail to complete the sexual cycle, which normally coincides with a peak of MTA1 expression. Thus, MTA1 functions in a developmental stage-specific manner. We determine the impact of 6mA on chromatin organization *in vitro* by reconstructing complete, full-length ciliate chromosomes harboring 6mA in native or ectopic positions. Using these synthetic chromosomes, we show that 6mA directly disfavors nucleosomes *in vitro* in a local, quantitative manner, independent of DNA sequence. Furthermore, the chromatin remodeler ACF can overcome this effect. Our study identifies a novel MT-A70 protein necessary for eukaryotic 6mA methylation and defines the impact of 6mA on chromatin organization using epigenetically defined synthetic chromosomes.

**Highlights:**

- The MT-A70 protein MTA1 mediates DNA N6-adenine methylation in *Oxytricha*
- MTA1 mutants exhibit subtle changes in nucleosome organization and transcription *in vivo*
- 6mA directly disfavors nucleosome occupancy in natural and synthetic chromosomes *in vitro*
- *De novo* synthesis of complete, epigenetically defined *Oxytricha* chromosomes

## Introduction

Covalent modifications on DNA have long been recognized as a hallmark of epigenetic regulation. DNA N6-methyladenine (6mA) has recently come under scrutiny in eukaryotic systems. Though previously thought to be primarily restricted to bacteria, where it functions as part of the restriction-modification (RM) system to protect host genomes from foreign elements (Bickle and Krüger, 1993), recent studies suggest its presence in plants and animals, with proposed roles in retrotransposon or gene regulation, transgenerational epigenetic inheritance, and chromatin organization (Luo et al., 2015). 6mA exists at low levels in *Arabidopsis thaliana* (0.006%-0.138% 6mA / dA), rice (0.2%), *C. elegans* (0.01–0.4%), *Drosophila* (0.001-0.07%), *Xenopus laevis* (0.00009%), mouse ES cells (0.0006 – 0.007%), human cells (Greer et al., 2015; Koziol et al., 2015; Liang et al., 2018; Wu et al., 2016; Xiao et al., 2018; Zhang et al., 2015; Zhou et al., 2018) and the mouse brain (Yao et al., 2017), although it accumulates in abundance (0.1–0.2%) during vertebrate embryogenesis (Liu et al., 2016). Disruption of DMAD, a 6mA demethylase, in the *Drosophila* brain leads to the accumulation of 6mA and Polycomb-mediated silencing (Yao et al., 2018). The existence of 6mA in mammals remains a subject of debate. Quantitative LC-MS analysis of HeLa and mouse ES cells failed to detect 6mA above background levels, raising concerns that initial observations of this base in mammals may in part be due to bacterial contamination (Schiffers et al., 2017). A recent study, however, reported that loss of 6mA in human cells promotes tumor formation (Xiao et al., 2018), suggesting that 6mA is a biologically relevant epigenetic mark in human disease.

In contrast to metazoans, 6mA is abundant in various unicellular eukaryotes, including ciliates (0.18 – 2.5%) (Ammermann et al., 1981; Cummings et al., 1974; Gorovsky et al., 1973; Rae and Spear, 1978), and the green algae *Chlamydomonas* (0.3 – 0.5%) (Fu et al., 2015; Hattman et al., 1978). High levels of 6mA (up to 2.8%) were also recently reported in basal fungi (Mondo et al., 2017). Ciliates thus constitute ideal systems for probing the function of 6mA, given its abundance and the availability of genetic and biochemical tools. Indeed, ciliates have long served as powerful models for the study of chromatin modifications (Brownell et al., 1996; Liu et al., 2007; Strahl et al., 1999; Taverna et al., 2002; Wei et al., 1998). Ciliates possess two structurally and functionally distinct nuclei within each cell (Prescott, 1994; Yerlici and Landweber, 2014). In *Oxytricha trifallax*, the germline *micronucleus* is transcriptionally silent and contains ~100 megabase-sized chromosomes, the same order of magnitude as most well studied model organisms (Chen et al., 2014). In contrast, the somatic *macronucleus* is transcriptionally active, being the sole locus of Pol II-dependent RNA production in non-developing cells (Khurana et al., 2014). The *Oxytricha* macronuclear genome is extraordinarily fragmented, consisting of ~16,000 unique chromosomes with a mean length of ~3.2kb, most encoding a single gene. Chromosome ends are capped with a compact 36nt telomere consisting of 5’-(T_4_G_4_)-3’ repeats, bound cooperatively by a heterodimeric protein complex (Gottschling and Zakian, 1986; Horvath et al., 1998). Macronuclear chromatin yields a characteristic ~200bp ladder upon digestion with micrococcal nuclease, indicative of regularly spaced nucleosomes (Gottschling and Cech, 1984; Lawn et al., 1978; Wada and Spear, 1980). Yet it remains unknown how and where nucleosomes are organized within these miniature chromosomes, and if this in turn regulates (or is regulated by) 6mA deposition.

Nucleosomes limit the physical accessibility of DNA to *trans*-acting factors, and thus directly impact DNA-based transactions, such as transcription, DNA repair, and replication. The *in vitro* reconstitution of chromatin is a method central to understanding these processes at the molecular level. Currently, the most widely used DNA template for such experiments is the 147bp “601” nucleosome positioning sequence. It exhibits a ~374-fold higher affinity for histone octamers than native genomic sequences (Lowary and Widom, 1998; Thåström et al., 2004) allowing the preparation of consistent, defined nucleosome arrays *in vitro*. Yet the biological relevance of 601 DNA remains unclear, due to its unnaturally high affinity for histones. On the other hand, reconstituting chromatin from native genomic DNA is challenging because long chromosomes increase the propensity for non-specific aggregation during chromatin assembly (Kaplan et al., 2009). As a compromise, small genomic fragments are usually prepared via PCR amplification or mechanical shearing. However, such substrates are devoid of their cognate physical environment, as defined by the totality of the chromosome. Here, we circumvent these limitations by exploiting the naturally small chromosomes of *Oxytricha*. Building ‘designer’ chromosomes with defined molecular composition permits us to study DNA modifications such as 6mA in the context of fully native DNA.

Intriguingly, in green algae, basal yeast, and the ciliate *Tetrahymena*, 6mA is enriched in ApT dinucleotide motifs within nucleosome linker regions near promoters (Fu et al., 2015; Hattman et al., 1978; Karrer and VanNuland, 1999; Mondo et al., 2017; Pratt and Hattman, 1981; Wang et al., 2017). These previous data suggested a correlation between 6mA, chromatin organization and gene regulation. Yet the functional impact of 6mA on these processes is still unclear, due to the inability to perturb methylation levels. Several questions remain open, such as whether 6mA regulates transcription or is a by-product of this process. Does 6mA directly disfavor nucleosomes, and if so, is it a graded or binary effect? Do ATP-dependent chromatin remodelers – both integral components of chromatin *in vivo* – modulate this interaction? Here we show that 6mA is localized in nucleosome linker regions near transcription start sites (TSSs) in *Oxytricha trifallax*. 6mA primarily occurs within ApT dinucleotide motifs, similar to algae, basal yeasts, and *Tetrahymena*. Through biochemical and genetic approaches, we identify a divergent MT-A70 domain protein – named MTA1 – necessary for 6mA methylation in *Oxytricha*. Disruption of MTA1 leads to genome-wide depletion of 6mA and small but significant increases in the fuzziness of flanking nucleosomes *in vivo*. Surprisingly, nucleosome organization is otherwise largely preserved in *mta1* mutants. To dissect the impact of 6mA on nucleosome organization *in vitro*, we built epigenetically defined *Oxytricha* chromosomes with or without 6mA at positions identical to their *in vivo* configuration, or at ectopic locations. Using this array of synthetic chromosomes, we show that 6mA directly disfavors nucleosome occupancy in a quantitative, site-specific manner *in vitro*, independent of DNA sequence. These effects are significantly greater than that observed *in vivo* but can be overcome by ATP-dependent chromatin remodelers. Intriguingly, RNAseq analysis revealed two classes of genes with different transcriptional responses to 6mA loss in *mta1* mutants, depending on their initial methylation level. Mutants fail to complete the sexual cycle, suggesting that 6mA is necessary for developmental progression. Together, we identify a novel MT-A70 protein as a key mediator of 6mA methylation, and determine the contribution of 6mA to gene expression and nucleosome organization. Our work also demonstrates the use of *Oxytricha* chromosomes as a versatile platform for functional studies of epigenetic modifications.

## Results

### Epigenomic profiles of chromatin and transcription in *Oxytricha*

We generated genome-wide *in vivo* maps of nucleosome positioning, transcription, and 6mA in the macronuclei of asexually growing (vegetative) *Oxytricha trifallax* cells using MNase-seq, poly(A)+ RNA-seq, 5’-complete cDNA-seq, and single molecule real time (SMRT) sequencing (Figure 1). *Oxytricha* TSSs localize within 100bp of chromosome ends, upstream of a phased nucleosome array (Figure 1A). Strikingly, 6mA is enriched in three consecutive nucleosome depleted regions directly downstream of TSSs. Each region contains varying levels of 6mA (Figure 1B), with the +1/+2 nucleosome linker being most densely methylated (Table S1A). In general, highly transcribed chromosomes tend to bear more 6mA, suggesting a positive role of this DNA modification in gene regulation (Figure 1C). The majority of methylation marks are located within an ApT motif (Figures 1D and 1E). 6mA occurs on sense and antisense strands with approximately equal frequency, indicating that the underlying methylation machinery does not function strand-specifically.

**Figure 1.**
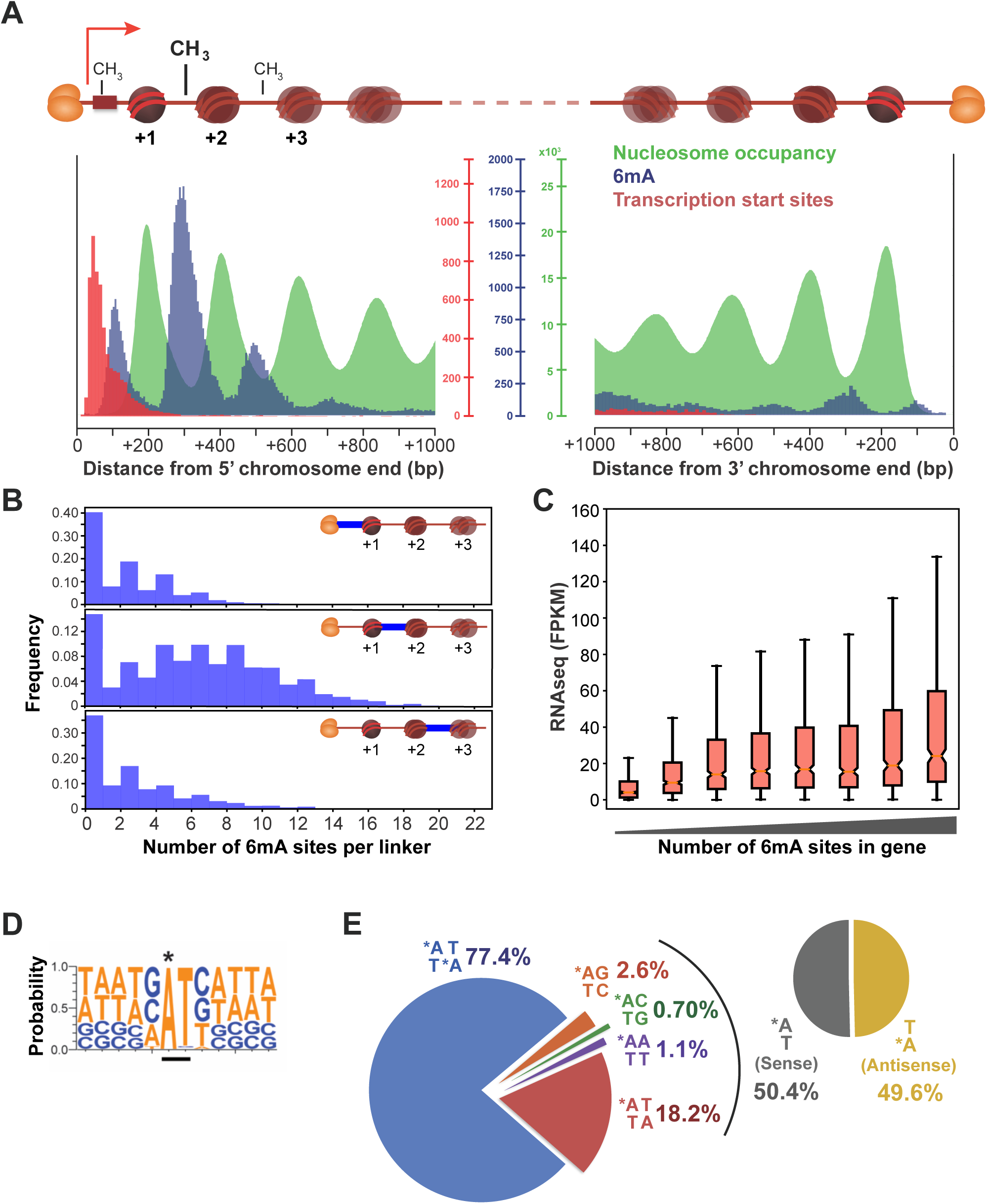
Epigenomic profiles of chromatin, transcription and DNA methylation in*Oxytricha* chromosomes. A. Meta-chromosome plots overlaying *in vivo* MNase-seq (nucleosome occupancy), SMRT-seq (6mA), and 5’-complete cDNA sequencing data (transcription start sites; TSSs) at *Oxytricha* macronuclear chromosome ends. Heterodimeric telomere end-binding protein complexes protect each end *in vivo* and are denoted as orange ovals. Horizontal red bar denotes the promoter. The 5’ chromosome end is designated as being proximal to TSSs. +1, +2, and +3 nucleosomes are labeled relative to the 5’ chromosome end. Vertical axis: “Nucleosome occupancy” denotes MNase-seq read coverage; “6mA” denotes total number of detected 6mA marks; “Transcription start sites” denotes total number of called TSSs.
B. Histograms of the total number of 6mA marks within each linker in *Oxytricha* chromosomes. Distinct linkers are depicted as horizontal bold blue lines.
C. Transcriptional activity is positively correlated with 6mA levels. RNAseq data are derived from poly(A)-enriched RNA. Genes are sorted into 8 groups, according to the total number of 6mA marks between 0 bp to 800 bp downstream of the TSS. FPKM = Fragments per Kilobase of transcript per Million mapped RNAseq reads. Notch in the boxplot denotes median, ends of boxplot denote first and third quartiles, upper whisker denotes third quantile + 1.5 × interquartile range, and lower whisker denotes data quartile 1 − 1.5 × interquartile range.
D. Composite analysis of 65,107 methylation sites reveals that 6mA (marked with **\***) occurs within an 5’-ApT-3’ dinucleotide motif.
E. Distribution of various 6mA dinucleotide motifs across the genome. Asterisk indicates DNA N6-methyladenine.

Using SMRT-seq base calls, the overall abundance of 6mA relative to (dA + 6mA) was calculated to be 0.78 – 1.04%. We independently verified the presence of 6mA in *Oxytricha* gDNA using quantitative LC-MS analysis with stable isotope-labeled internal nucleoside standards (Figure S1). This experiment yielded a 6mA / (dA + 6mA) ratio of 0.71%. Note that the calculation from SMRT-seq data is expected to be an overestimate because 6mA is scored at being present or absent at each site in the genome for this purpose. In actual fact, 6mA sites may be partially methylated (Figure S2A). Neither 6mA nor dA was detected from LC-MS analysis of *Oxytricha* culture media, arguing against spurious signal arising from contamination or overall technical handling. Our PacBio and LC-MS measurements of % 6mA are both similar to thin layer chromatography analysis of nucleotides (0.6 – 0.7%) from a distinct but closely related species, *Oxytricha fallax* (Rae and Spear, 1978).

### MTA1 is an MT-A70 protein necessary for 6mA methylation *in vivo*

To uncover the functions of 6mA *in vivo*, we set out to identify and disrupt putative 6mA methytransferases (MTases). The *Oxytricha* macronuclear genome encodes five genes belonging to the MT-A70 family (Table S2) (Iyer et al., 2016; Swart et al., 2013). Such genes commonly function as RNA m^6^A MTases in eukaryotes, having evolved from m.MunI-like MTases in bacterial restriction-modification systems (Iyer et al., 2016). An MT-A70 gene belonging to the METTL4 subclade, DAMT-1, is a putative 6mA methyltransferase in *C. elegans* (Greer et al., 2015). However, none of the *Oxytricha* MT-A70 genes clustered together with METTL4 on a phylogenetic tree (Figure 2A). The *Oxytricha* genome also contains homologs of a structurally distinct RNA m^6^A MTase, METTL16, which was reported to methylate U6 snRNA (Table S2) (Pendleton et al., 2017; Warda et al., 2017). Another candidate, N6AMT1 – which does not contain an MT-A70 domain – was recently found to mediate DNA 6mA methylation in human cells (Xiao et al., 2018). An N6AMT1 homolog is also present in the *Oxytricha* genome (Table S2). The large number of candidate methyltransferases rendered it impractical to test gene function, one at a time or in combination. To identify the ciliate 6mA MTase, we undertook a biochemical approach by fractionating nuclear extracts and identifying candidate proteins that co-purified with DNA methylase activity. The organism of choice for this experiment was *Tetrahymena thermophila*, a ciliate that divides at a significantly faster rate than *Oxytricha* (~2 hours vs 18 hours); (Cassidy-Hanley, 2012; Laughlin et al., 1983). The faster growth time of *Tetrahymena* rendered it feasible to culture large amounts of cells for nuclear extract preparation. We assessed 6mA positions in *Tetrahymena* gDNA using SMRT-seq and observed a similar genomic localization as *Oxytricha*, enriched in a consensus ApT motif within nucleosome linker regions near transcription start sites (Figure S3). We independently verified that 6mA is present in *Tetrahymena* gDNA through quantitative LC-MS (Figure S1), in agreement with earlier reports (Gorovsky et al., 1973; Pratt and Hattman, 1981). We thus reasoned that the enzymatic machinery responsible for 6mA deposition is conserved between *Tetrahymena* and *Oxytricha*, and that *Tetrahymena* could serve as a tractable biochemical system for identifying the ciliate 6mA MTase.

**Figure 2.**
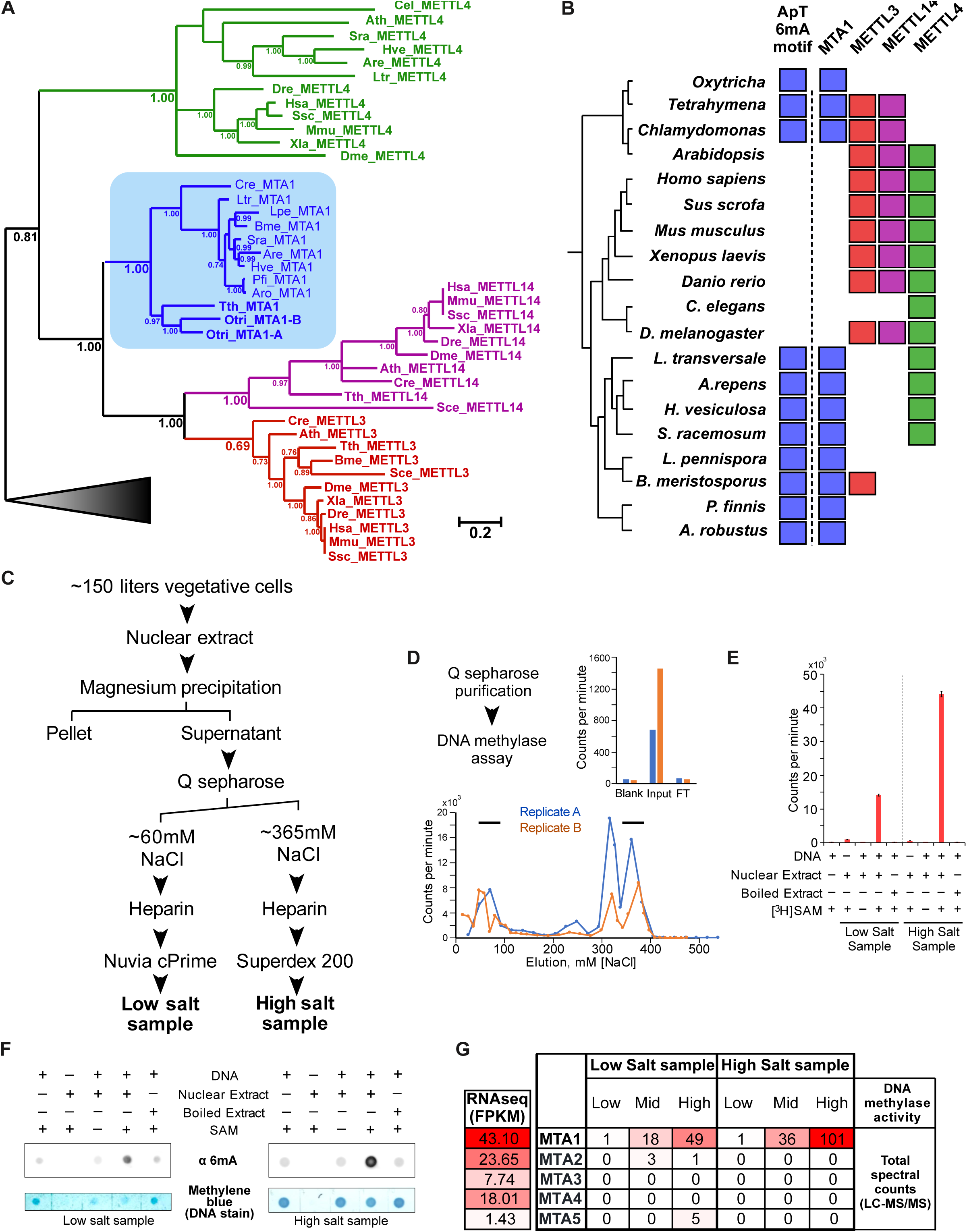
Purification and identification of the ciliate 6mA methyltransferase. A. Bayesian phylogenetic tree depicting relationship of MTA1 to other MT-A70 proteins. Posterior probabilities greater than > 0.65 are shown. Gray triangle represents outgroup of bacterial sequences. Abbreviations: Cel: *Caenorhabditis elegans*; Ath: *Arabidopsis thaliana*; Sra: *Syncephalastrum racemosum*; Hve: *Hesseltinella vesiculosa*; Are: *Absidia repens*; Dre: *Danio rerio*; Has: *Homo sapiens*; Ssc: *Sus scrofa*; Mmu: *Mus musculus*; Xla: *Xenopus laevis*; Dme: *Drosophila melanogaster*; Cre: *Chlamydomonas reinhardtii*; Ltr: *Lobosporangium transversale*; Lpe: *Linderina pennispora*; Bme: *Basidiobolus meristosporus*; Pfi: *Piromyces finnis*; Aro: *Anaeromyces robustus*; Tth: *Tetrahymena thermophila*; Otri: *Oxytricha trifallax*.
B. Phylogenetic distribution of the occurrence of ApT 6mA motifs and various MT-A70 protein families, including MTA1. Filled square denotes the presence of a particular MT-A70 family member in a taxon. The basal yeast clade is comprised of *L. transversale*, *A.repens*, *H. vesiculosa*, *S. racemosum*, *L. pennispora*, *B. meristosporus*, *P. finnis*, and *A. robustus*.
C. Experimental scheme depicting the partial purification of DNA methyltransferase activity from *Tetrahymena* nuclear extracts. Individual fractions from each purification step were assayed for DNA methyltransferase activity as described in panel E. Fractions that elute from the Nuvia cPrime and Superdex 200 columns were further analyzed by mass spectrometry to identify candidate proteins that co-elute with DNA methyltransferase activity. Mass spectrometry data are described in panel G.
D. Fractionation of nuclear extracts on Q sepharose column results in two distinct peaks of DNA methyltransferase activity, denoted as “Low Salt sample” and “High Salt sample” by black horizontal bars. FT denotes flow-through from column. The DNA methyltransferase assay involves mixing DNA substrate, nuclear extract, and radiolabeled SAM cofactor. Incorporation of radioactivity into the DNA substrate is measured using scintillation counting. The salt concentration at which individual fractions elute from the column is plotted against DNA methyltransferase activity of each fraction (counts per minute). Inset shows DNA methyltransferase activity of the input nuclear extract, flowthrough from the Q sepharose column, and blank control (nuclear extract buffer). Orange and blue plots denote replicates derived from independent preparations of nuclear extract.
E. DNA methyltransferase assay showing that the activity is heat-sensitive and requires DNA and radiolabeled SAM. Error bars represent s.e.m. (n = 3).
F. Dot blot showing that nuclear extracts mediate 6mA methylation. Note that the low salt sample has substantial DNase activity, resulting in a lower amount of DNA available for dot blot analysis. DNA substrate, nuclear extract, and SAM cofactor were mixed as in panels D and E. The DNA was subsequently purified and used for dot blot analysis.
G. RNAseq gene expression and protein abundance of *Tetrahymena* MT-A70 genes in partially purified nuclear extracts. UniProt IDs for each gene is listed in Table S2. RNAseq data are obtained from (Xiong et al., 2012). Both RNAseq and protein mass spectrometry data are generated from the same physiological state (log-phase vegetative *Tetrahymena* cells). FPKM = Fragments per Kilobase of transcript per Million mapped RNAseq reads. A higher FPKM value indicates a greater normalized RNAseq read count and thus higher gene expression. “Low”, “Mid”, and “High” DNA methylase activity correspond to distinct fractions eluting from the Nuvia cPrime and Superdex 200 columns in panel C. These fractions exhibit qualitatively different DNA methylase activity as measured using scintillation counting. Mass spectrometry analysis was subsequently performed on these fractions to identify candidate proteins whose abundance correlates with activity. “Total spectrum counts” is a semi-quantitative measure of protein abundance, representing the total number of LC-MS/MS fragmentation spectra that match peptides from a target protein. The extent of shading in each box is proportional to RNAseq or spectral count values.

We prepared nuclear extracts from log-phase *Tetrahymena* cells, since 6mA could be readily detected at this developmental stage through quantitative mass spectrometry and PacBio sequencing (Figures S1 and S3). Nuclear extracts were incubated with radiolabeled S-adenosyl-L-methionine and PCR-amplified DNA substrate to assay for DNA methylase activity. Passage of the nuclear extract through an anion exchange column resulted in the elution of two distinct peaks of DNA methylase activity, both of which were heat sensitive (Figure 2C – 2E). Western blot analysis confirmed that both peaks of activity mediate methylation on 6mA (Figure 2F). The resulting fractions were further purified and subjected to mass spectrometry. One MT-A70 protein – termed MTA1 – was detected at higher abundance in fractions with high DNA methylase activity. MTA1 received the large majority of peptide matches, relative to all other MT-A70 genes encoded by the *Oxytricha* genome (Figures 2G and S4). MTA1 was detected in both peaks of DNA methylase activity that eluted from the anion exchange column, suggesting that the protein may form distinct active complexes *in vivo*. Curiously, although poly(A)-selected RNA transcripts are present from all MT-A70 genes (Figure 2G), almost all peptides in fractions with high DNA methylase activity corresponded to MTA1. No peptides corresponding to METTL16 or N6AMT1 homologs were detected in any fraction. We then examined the level of MTA1 poly(A)+ RNA transcripts at various developmental stages of *Tetrahymena* and observed a > 5-fold increase in expression early in the sexual cycle, relative to log-phase vegetative growth (Figure S5). We compared this to recently published 6mA immunofluorescence data from developmentally staged *Tetrahymena* cells (Wang et al., 2017). Strikingly, the peak in MTA1 expression early in sexual development coincides with a sharp increase in nuclear 6mA staining, indicating that the accumulation of MTA1 and 6mA correlate *in vivo*.

We next investigated the phylogenetic relationship of MTA1 to other eukaryotic MT-A70 domain-containing proteins. Two widely studied mammalian MT-A70 proteins – METTL3 and METTL14 (Ime4 and Kar4 in yeast) – form a heterodimeric complex that is responsible for m^6^A methylation in mRNA. METTL3 is the catalytically active subunit, while METTL14 functions as an RNA-binding scaffold protein (Śledź and Jinek, 2016; Wang et al., 2016a, 2016b). MTA1 lies outside of the monophyletic clades formed by mammalian METTL3 and METTL14, and *C. elegans* DAMT-1 (METTL4) (Figure 2A). Thus, MTA1 is a diverged MT-A70 family member that is phylogenetically distinct from all previously studied RNA and DNA N6-methyladenine MTases. We then asked whether MTA1 is also found in other eukaryotes with a similar occurrence of 6mA in ApT motifs as *Tetrahymena*. We queried the genomes of *Oxytricha*, green algae, and eight basal yeast species, all of which exhibit this distinct methylation pattern (as evidenced from Figure 1, Figure S3, (Fu et al., 2015), and (Mondo et al., 2017)). For all of these taxa, we could recover MT-A70 homologs that grouped monophyletically with MTA1 (Figure 2B). On the other hand, MT-A70 homologs from multicellular eukaryotes, including *Arabidopsis*, *C. elegans*, *Drosophila*, and mammals, grouped exclusively with METTL3, METTL14, and METTL4 lineages, but not MTA1. None of these latter genomes exhibit a consensus ApT dinucleotide methylation motif for 6mA (Greer et al., 2015; Koziol et al., 2015; Liang et al., 2018; Liu et al., 2016; Wu et al., 2016; Xiao et al., 2018; Zhang et al., 2015). We note that the absence of an ApT dinucleotide motif is based on data from a limited number of cell types, developmental stages, and culture conditions tested in these studies. Nonetheless, within the scope of currently available data, the presence of MTA1 correlates with the distinctive genomic localization of 6mA within ApT motifs.

To directly test the role of MTA1 in mediating 6mA methylation, we disrupted *Oxytricha* MTA1 (paralog MTA1-A, Figure 2A) by inserting an ectopic DNA sequence 49 bp downstream of the start codon. This resulted in a frameshift mutation and loss of the C-terminal MTase domain (Figure 3A). RNAseq analysis confirmed the presence of the ectopic insertion in *mta1* mutant transcripts but not wild type controls (Figure S6). *Oxytricha* has two MTA1 paralogs, but we focused on MTA1-A because MTA1-B RNA transcripts are not detected in vegetative *Oxytricha* cells (Swart et al., 2013), where we profiled 6mA locations through SMRT-seq. Dot blot analysis confirmed a significant reduction in bulk 6mA levels in mutant lines (Figure 3B). We then examined 6mA positions at high resolution using SMRT-seq to understand how the DNA methylation landscape is altered in *mta1* mutants. Strikingly, these mutants exhibit genome-wide loss of 6mA, with complete abolishment of the dimethylated ApT motif, and reduction in frequency of all other methylated dinucleotide motifs (Figures 3C-E). These findings are consistent across all biological replicates and are robust to wide variation in SMRT-seq parameters for calling 6mA modifications (Figure S2B-S2D). It cannot be attributed to variation in sequencing coverage between wild type and mutant lines. The Inter Pulse Duration ratio (degree of polymerase slowing during PacBio sequencing due to presence of a modified base) and estimated fractional methylation also decreased significantly at called 6mA sites in *mta1* mutants (p < 2.2 x 10^−16^, Wilcoxon rank-sum test) (Figure S2A). MTA1 is therefore necessary for a significant proportion of *in vivo* 6mA methylation events in *Oxytricha*.

**Figure 3.**
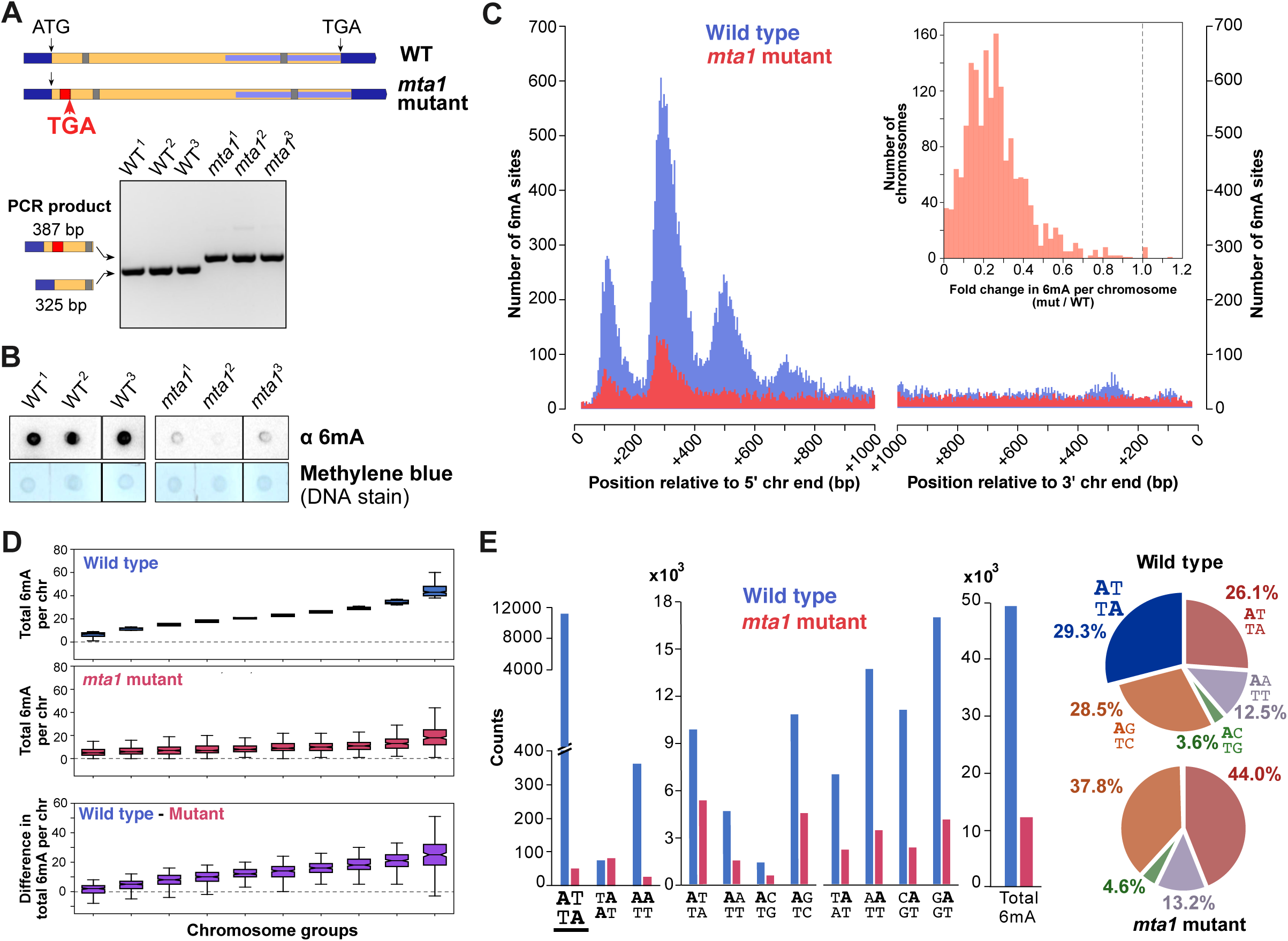
Genome-wide loss of 6mA in *mta1* mutants. A. Schematic depicting the disruption of *Oxytricha* MTA1 open reading frame. Flanking dark blue bars: 5’ and 3’ UTR; yellow = open reading frame; red = retention of 62 bp ectopic DNA segment that is normally deleted during sexual development. This leads to a frameshift mutation and premature translation termination; grey bar = intron; internal light blue bar = annotated MT-A70 domain; “ATG” = start codon; “TGA” = stop codon. Agarose gel analysis shows PCR confirmation of ectopic DNA retention within the MTA1 open reading frame in *mta1* mutant lines.
B. Dot blot of RNase-treated genomic DNA showing decrease of 6mA in *mta1* mutants.
C. Histogram of 6mA counts near 5’ and 3’ Oxytricha chromosome ends, showing drastic decrease in 6mA levels in *mta1* mutants. Inset depicts histogram of fold change in total 6mA in each chromosome, between mutant and wild type cell lines.
D. Chromosomes are divided into 10 equal groups, ranked according to total 6mA in each chromosome in wild type cells (blue boxplots). For each group, the total 6mA per chromosome in mutants, and the difference in total 6mA per chromosome is plotted below. Notch in the boxplot denotes median, ends of boxplot denote first and third quartiles, upper whisker denotes third quantile + 1.5 × interquartile range, and lower whisker denotes data quartile 1 − 1.5 × interquartile range.
E. Dinucleotide motif distribution in wild type and *mta1* mutants. Note the loss of ApT dimethylated motif in *mta1* mutants (underlined). Bar charts represent absolute counts of motifs, while pie charts show relative proportion of motifs.

What are the phenotypic consequences of 6mA loss *in vivo*? It has been proposed that DNA methylation – including 6mA and cytosine methylation – is involved in nucleosome organization (Fu et al., 2015; Huff and Zilberman, 2014). We thus asked whether nucleosome organization is altered in *mta1* mutants. We quantified nucleosome “fuzziness”, defined as the standard deviation of MNase-seq read locations surrounding the called nucleosome peak (Lai and Pugh, 2017; Mavrich et al., 2008). A poorly positioned nucleosome consists of a shallow and wide peak of MNase-seq reads, manifested by a high fuzziness score. In *mta1* mutants, the nucleosomes that experience large changes in flanking 6mA do increase significantly in fuzziness (Figure S7A and S7D). Such nucleosomes also exhibit changes in occupancy that are consistent with an increase in fuzziness (Figures S7A and S7E). These results are robust to variation in MNase digestion (Figure S8C and S8D). On the other hand, nucleosome linkers do not change in length or occupancy, even though 6mA is lost from these regions (Figures S7B,C,F,G and S9). Note that an increase in nucleosome fuzziness does not necessitate an increase in adjacent linker occupancy, as these metrics are calculated from non-overlapping windows (see Figure S7C legend for explanation). We conclude that 6mA exerts subtle effects on nucleosome organization *in vivo*.

### 6mA disfavors nucleosome occupancy across the genome *in vitro* but not *in vivo*

Multiple factors, including 6mA, DNA sequence, and chromatin remodeling complexes, may collectively contribute to nucleosome organization *in vivo*. The effect of 6mA could therefore be masked by these elements. We next sought to determine whether 6mA directly impacts nucleosome organization. To this end, we assembled chromatin *in vitro* using *Oxytricha* gDNA, which contains cognate 6mA. To obtain a matched negative control lacking DNA methylation, we exploited the naturally fragmented architecture of the *Oxytricha* macronuclear genome to amplify complete chromosomes using PCR. This erases all 6mA marks, while fully preserving DNA sequence and physical linkage within each chromosome. We selected 98 unique chromosomes that collectively reflect overall genome properties, including AT content, chromosome length, and transcriptional activity (Table S1B). Only high copy number chromosomes were selected to ensure high SMRT-seq coverage and thus confident identification of 6mA marks. Full-length chromosomes were individually PCR-amplified from genomic DNA, resulting in the collective erasure of 2,344 6mA marks (Figure 4A). Each chromosome was purified and subsequently mixed together in stoichiometric ratios to obtain a “mini-genome” (Figure 4B). Native genomic DNA (containing 6mA) and amplified mini-genome DNA (lacking 6mA) were each assembled into chromatin *in vitro* using *Xenopus* or *Oxytricha* histone octamers (Figures S10, S11A and S11B) and analyzed using MNase-seq. We computed nucleosome occupancy from the native genome and mini-genome samples across 199,795 overlapping DNA windows, spanning all base-pairs in the 98 chromosomes. This allowed the direct comparison of nucleosome occupancy in each window of identical DNA sequence, with and without 6mA (Figures 4C and 4D). Windows exhibit lower nucleosome occupancy with increasing 6mA, confirming the quantitative nature of this effect. Furthermore, similar trends were observed for both native *Oxytricha* and recombinant *Xenopus* histones, suggesting that the effects of 6mA on nucleosome organization arise mainly from intrinsic features of the histone octamer rather than from species-specific variants (Figure 4C and 4D). These results are also robust to the extent of MNase digestion of reconstituted chromatin (Figure S8A).

**Figure 4.**
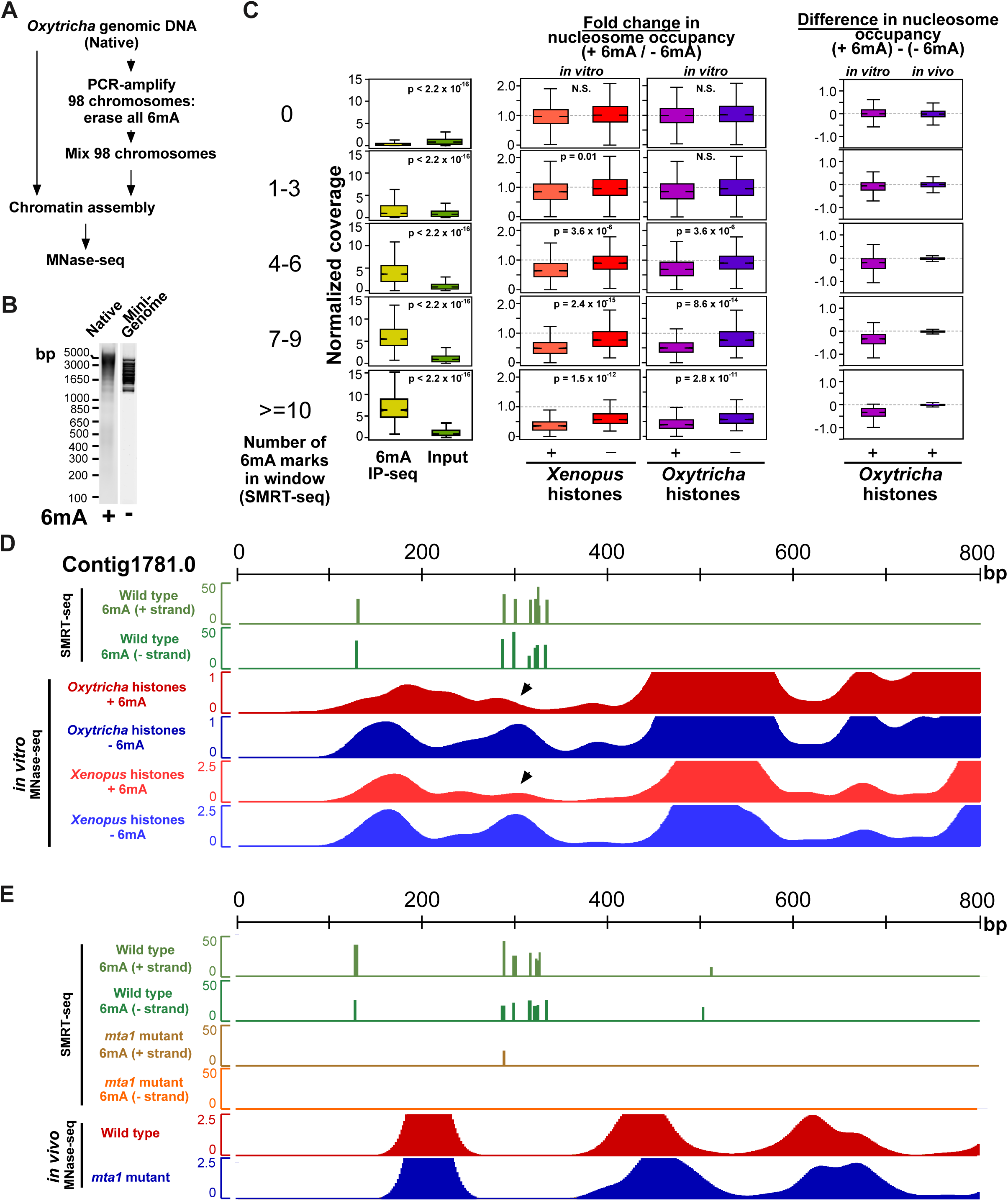
Effects of 6mA on nucleosome organization *in vitro* and *in vivo*. A. Experimental workflow for the generation of mini-genome DNA from native genomic DNA, and subsequent analysis by MNase-seq.
B. Agarose gel analysis of *Oxytricha* gDNA (‘Native’) and mini-genome DNA before chromatin assembly.
C. Methylated regions in the genome exhibit lower nucleosome occupancy *in vitro* but not *in vivo*. MNase-seq (nucleosome occupancy) and 6mA IP-seq coverage were calculated within overlapping 51bp windows across the 98 assayed chromosomes. Windows were binned according to the number of 6mA residues within. The *in vitro* MNase-seq coverage from chromatinized native gDNA (“+” 6mA) was divided by the corresponding coverage from chromatinized mini-genome DNA (“-“ 6mA) to obtain the fold change in nucleosome occupancy in each window (“+” histones). A subtraction was performed on these datasets to obtain the difference in nucleosome occupancy *in vitro*. Identical DNA sequences were compared for each calculation. Naked native gDNA and mini-genome DNA were also MNase-digested, sequenced and analyzed in the same manner, to control for MNase sequence preferences (“-“ histones). Nucleosome occupancy *in vivo* corresponds to MNase-seq coverage from wild type and *mta1* mutant cells. *P*-values were calculated using a two-sample unequal variance t-test. N.S denotes “non-significant”, with p > 0.05.
D. Tracks of 6mA distribution and MNase-seq coverage reveal a reduction in nucleosome occupancy at methylated loci *in vitro* (black arrowheads). 6mA “+” strand and “-“ strand tracks refer to SMRT-seq base calls from wild type genomic DNA. For the *in vitro* MNase-seq tracks, “+ 6mA” refers to chromatin assembled on *Oxytricha* gDNA, while “-6mA” denotes chromatin assembled on mini-genome DNA. Similar trends are observed for both *Oxytricha* and *Xenopus* histones. Vertical axis for SMRT-seq data denotes confidence score [-10 log(p-value)] of detection of the methylation event, while that for *in vitro* MNase-seq data denotes normalized read coverage.
E. Tracks of 6mA distribution and MNase-seq coverage *in vivo* reveal no change in nucleosome occupancy in linker regions despite loss of 6mA in *mta1* mutants. Vertical axis for SMRT-seq tracks denote confidence score [-10 log(p-value)] of detection of the methylation event, while that for *in vivo* MNase-seq tracks denote normalized read coverage.

We then directly compared the impact of 6mA on nucleosome occupancy *in vitro* and *in vivo*. Loss of 6mA *in vitro* is achieved by mini-genome construction, while loss *in vivo* is achieved by the *mta1* mutation. For each overlapping DNA window, we calculated the difference in nucleosome occupancy: 1) between native genome and mini-genome DNA *in vitro*, and 2) between wild type and *mta1* mutants *in vivo* (Figure 4C). Nucleosome occupancy is indeed lower in the presence of 6mA methylation *in vitro* (Figures 4C and 4D). In contrast, no change in nucleosome occupancy is observed *in vivo* (Figures 4C and 4E). This result is consistent with our earlier analysis of linker occupancy in *mta1* mutants (Figure S7C and S7G). We note that highly methylated DNA windows show greater change in 6mA relative to *mta1* mutants (Figure 3D). Yet, these windows do not change in nucleosome occupancy *in vivo*. We conclude that 6mA methylation locally disfavors nucleosome occupancy *in vitro*, but that this intrinsic effect can be overcome by endogenous chromatin factors *in vivo*.

### Modular synthesis of epigenetically defined chromosomes

The above experiments used kinetic signatures from SMRT-seq data to infer the presence of 6mA marks in genomic DNA. We next sought to confirm that 6mA is directly responsible for disfavoring nucleosomes *in vitro*, and to understand how this effect could be overcome by cellular factors. We built synthetic, chemically defined chromosomes *in vitro* with fully native DNA sequence, containing 6mA marks installed via oligonucleotide synthesis. Importantly, 6mA was placed at all locations identified by SMRT-seq *in vivo*. These chromosomes were constructed *de novo* via ligation of individual DNA building blocks, themselves generated in large quantities through PCR. 6mA was introduced into oligonucleotides before ligation with flanking DNA building blocks, thus localizing the epigenetic mark to desired sites in the chromosome. We developed a streamlined chromosome synthesis scheme involving consecutive restriction enzyme digestion, ligation, and size selection steps (see Methods and Figure S12). These epigenetically defined chromosomes were then used to dissect the effect of 6mA on nucleosome occupancy. The representative chromosome, *Contig1781.0*, is 1.3kb, contains a clearly defined TSS, and encodes a single highly transcribed gene with a predicted RING finger domain. The length and gene structure are characteristic of typical *Oxytricha* chromosomes (Figure 5A). We independently validated the location of 6mA *in vivo* by sequencing chromosomal DNA immunoprecipitated with an anti-6mA antibody (Figure 5A).

**Figure 5.**
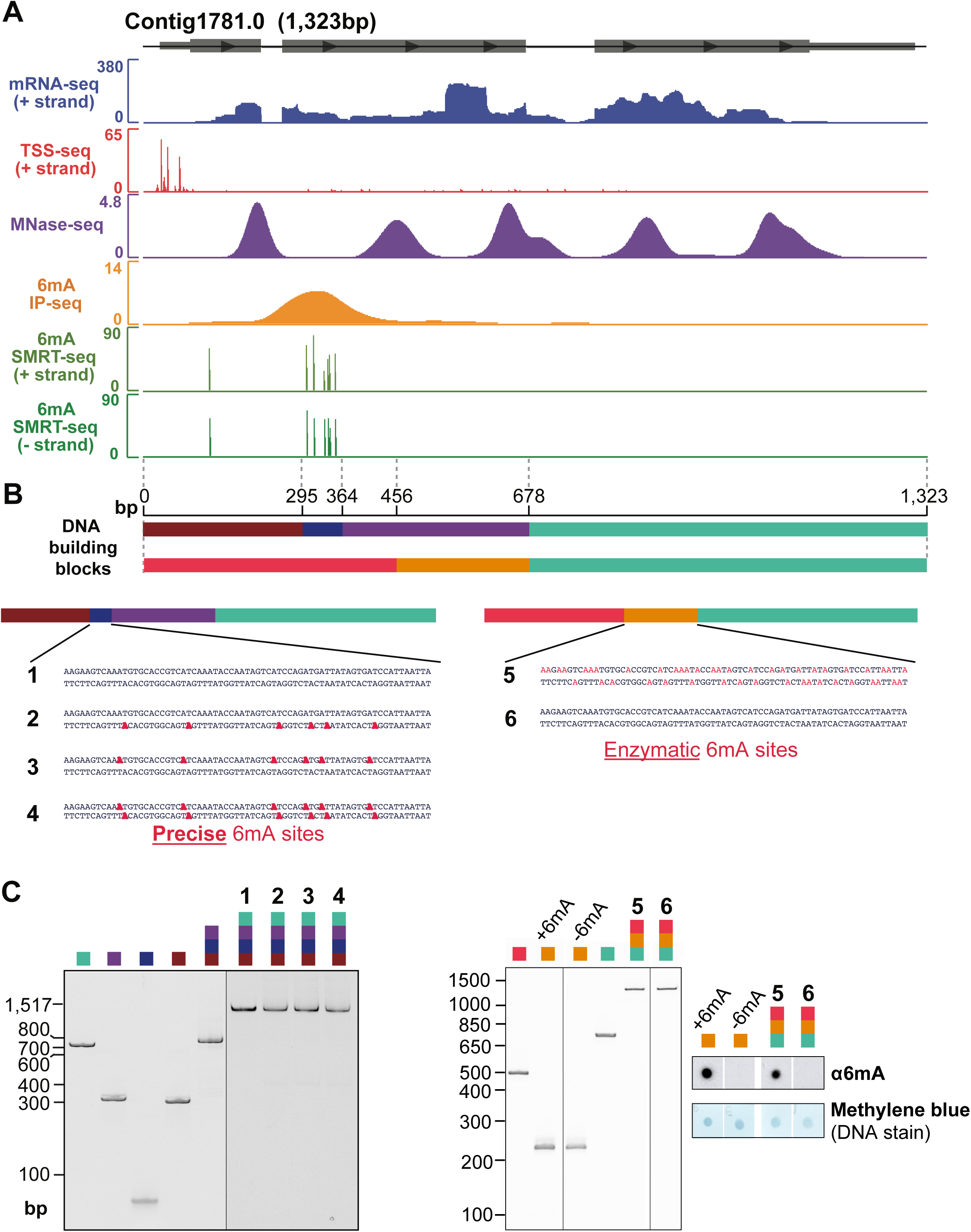
Modular synthesis of full-length *Oxytricha* chromosomes. A. Chromatin and transcriptional features used to select, design and synthesize a full-length chromosome (Contig1781.0). Horizontal grey boxes represent exons for the single annotated gene within this chromosome, with arrows denoting its orientation. All data tracks represent normalized sequencing coverage except for SMRT-seq, which represents the confidence score [-10 log(p-value)] of detection of each methylated base.
B. Schematic of building blocks used to construct synthetic *Oxytricha* chromosomes. Different colors denote separate DNA building blocks ligated to form the 1,323bp full-length chromosome. Each building block is drawn to scale. Two sets of building blocks are used to construct chromosomes 1-4 and 5-6, respectively. Precise 6mA sites (bold red) represent cognate 6mA positions revealed by SMRT-seq in native genomic DNA. These are introduced via oligonucleotide synthesis. For chromosome 5, enzymatic 6mA sites (non-bold red) represent possible locations installed by a bacterial 6mA methyltransferase.
C. Native polyacrylamide gel analysis of building blocks and purified synthetic chromosomes. Anti-6mA dot blot analysis confirms DNA methylation on synthetic chromosome.

Four chromosome variants were synthesized, with cognate 6mA sites on neither, one, or both DNA strands (chromosomes 1–4 in Figures 5B, 5C, and S12A). Each chromosome was cloned and sequenced to verify the accuracy of construction (Figure S12C). With these chromosomal DNA templates in hand, we investigated the impact of 6mA on nucleosome occupancy. Chromatin was assembled by salt dialysis with either *Oxytricha* or *Xenopus* nucleosomes and subsequently digested with MNase to obtain mononucleosomal DNA (Figures 6A and S11C). Tiling qPCR was used to quantify nucleosome occupancy at ~50bp increments along the entire length of the synthetic chromosome (Figure 6B). The fully methylated locus exhibits a ~46% reduction in nucleosome occupancy relative to the unmethylated variant, while hemimethylated chromosomes containing half the number of 6mA marks showed intermediate nucleosome occupancy at the corresponding region (Figures 6B). The reduction in nucleosome occupancy was confined to the methylated region, and not observed across the rest of the chromosome. Similar trends were observed when chromatin was assembled using the NAP1 histone chaperone (Figure S13B, top panel), indicating that this effect is not an artifact of the salt dialysis method. We therefore conclude that 6mA directly disfavors nucleosome occupancy in a local, quantitative manner *in vitro*.

**Figure 6.**
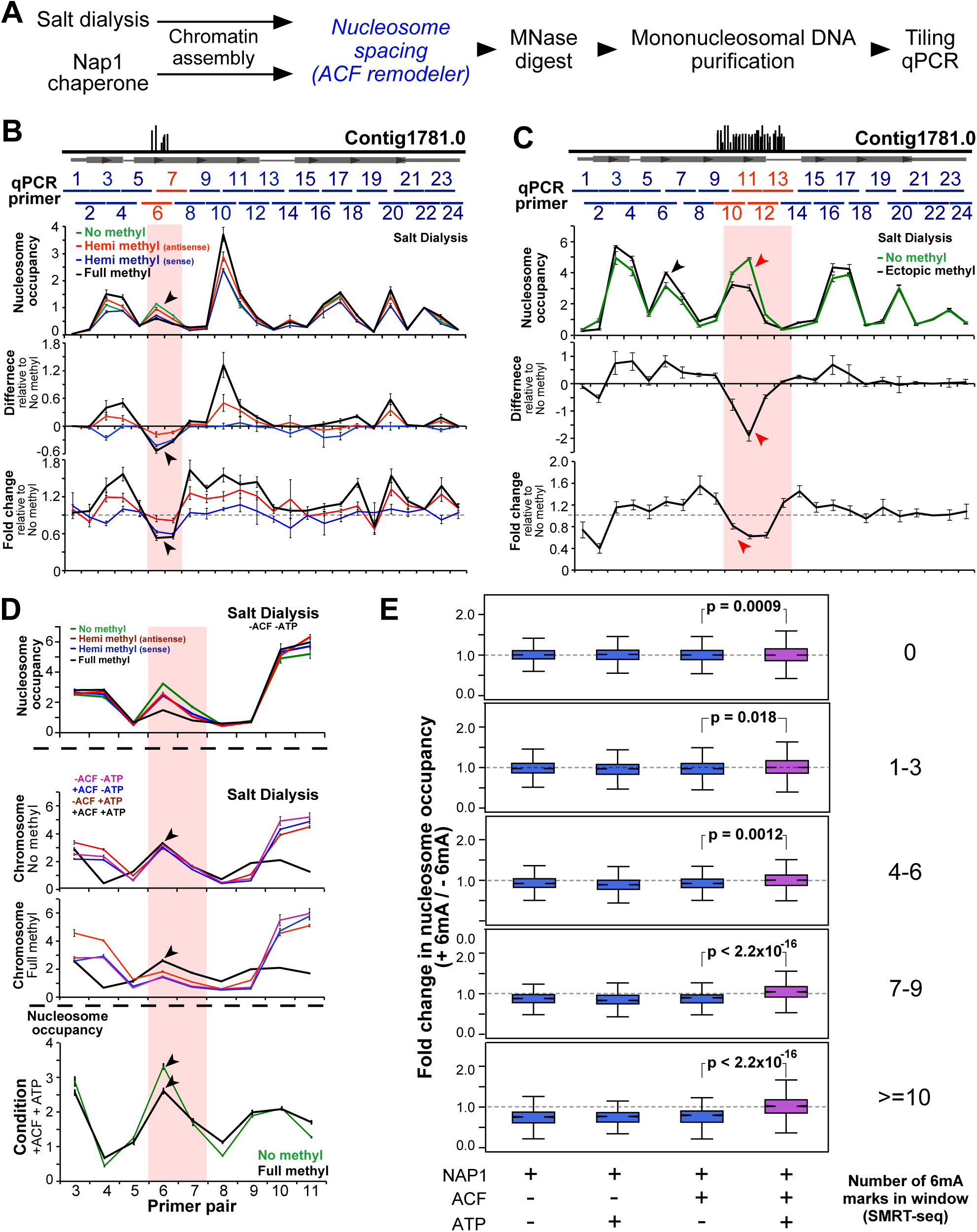
Quantitative modulation of nucleosome occupancy by 6mA in synthetic chromosomes. A. Experimental workflow. Chromatin is assembled using either salt dialysis or the NAP1 histone chaperone. Italicized blue steps are selectively included.
B. Tiling qPCR analysis of nucleosome occupancy in the synthetic chromosome Contig1781.0, with cognate 6mA sites. Horizontal grey box represents the annotated gene within Contig1781.0, with vertical grey lines depicting 6mA positions as obtained from Figure 5A. Horizontal blue bars span ~100bp regions amplified by qPCR primer pairs. Red horizontal lines and vertical bars represent the region containing 6mA. ‘Hemi methyl’ chromosomes contain 6mA on the antisense and sense strands, respectively, while the ‘Full methyl’ chromosome has 6mA on both strands. Nucleosome occupancy represents normalized qPCR signal at each locus (see Methods). For each qPCR locus, “difference” = nucleosome occupancy in methylated chromosome – nucleosome occupancy in “no methyl chromosome”; “fold change” = nucleosome occupancy in methylated chromosome / nucleosome occupancy in no methyl chromosome. Black arrowheads denote the decrease in nucleosome occupancy specifically at the 6mA cluster.
C. Tiling qPCR analysis of nucleosome occupancy in synthetic chromosome Contig1781.0 with ectopic 6mA sites. Vertical black lines illustrate possible 6mA sites installed enzymatically. Red arrowheads denote the decrease in nucleosome occupancy in the ectopically methylated region, while black arrowheads denote the position of cognate 6mA sites (not in this construct).
D. The chromatin remodeler ACF can partially overcome the effects of 6mA on nucleosome organization in an ATP-dependent manner. Chromatin is assembled by salt dialysis on synthetic chromosomes from panel B, and subsequently incubated with ACF and/or ATP. ACF equalizes nucleosome occupancy between the 6mA cluster and flanking regions in the presence of ATP (black line), resulting in a relative increase in nucleosome occupancy across the methylated region. Nucleosome occupancy at the methylated region is not restored to the same level as the unmethylated control in the presence of ACF and ATP (black arrowheads). **(E)** Chromatin is assembled on native gDNA (“+” 6mA) and mini-genome DNA (“-“ 6mA) using NAP1 in the presence of ACF and/or ATP, and subsequently analyzed using MNase-seq. Sliding window fold change in nucleosome occupancy between native gDNA and mini-genome DNA is calculated as in Figure 4C. ACF acts in an ATP-dependent manner to restore nucleosome occupancy in methylated DNA windows. *P*-values were calculated using a two-sample unequal variance t-test.

If 6mA impacts nucleosome occupancy regardless of the underlying DNA sequence, this effect should also be observed when 6mA is moved to a non-native site. To test this, we enzymatically installed 6mA at an ectopic location within the synthetic chromosome (chromosome 5 in Figures 5B, 5C and S12B). This region exhibits high *in vitro* nucleosome occupancy in the absence of 6mA (Figure 6C). The full-length chromosome was built by DNA ligation (Figures 5C and S12B) and subsequently assembled into chromatin by salt dialysis. Tiling qPCR analysis of mononucleosomal DNA showed a significant decrease in nucleosome occupancy at the ectopically methylated region, but not where native 6mA sites were originally present (Figure 6C). These data indicate that the impact of 6mA on nucleosome occupancy is qualitatively independent of DNA sequence. Future studies are required to determine whether specific DNA sequence features quantitatively influence this effect.

### Chromatin remodelers restore nucleosome occupancy over 6mA sites

Nucleosome occupancy *in vivo* is influenced not only by DNA sequences, but also by *trans*-acting factors. ATP-dependent chromatin remodeling factors modulate nucleosome organization and help establish canonical nucleosome patterns near TSSs (Struhl and Segal, 2013). We used synthetic, methylated chromosomes to test how the well-studied chromatin remodeler ACF responds to 6mA in native DNA. ACF generates regularly spaced nucleosome arrays *in vitro* and *in vivo* (Clapier and Cairns, 2009; Ito et al., 1997). Its catalytic subunit ISWI is conserved across eukaryotes, including *Oxytricha* and *Tetrahymena* (Figure S14). Synthetic chromosomes were assembled into chromatin by salt dialysis as before, then incubated with ACF in the presence of ATP (Figure S11D). We find that ACF partially – but not completely – restores nucleosome occupancy over the methylated locus in an ATP-dependent manner (Figure 6D). This effect is observed when ACF was added to chromatin assembled by salt dialysis or the NAP1 histone chaperone (Figure 6D and S13B). We then asked whether the impact of ACF on nucleosome organization is also observed in other methylated regions across the genome. To this end, we assembled native genomic DNA (containing 6mA) and mini-genome DNA (lacking 6mA) into nucleosomes using NAP1 and ACF (Figure S11E; An and Roeder, 2003; Fyodorov and Kadonaga, 2003). Mononucleosomal DNA was purified and subjected to deep sequencing. Indeed, ACF restores nucleosome occupancy over methylated regions in the genome in an ATP-dependent manner (Figure 6E). This result is robust to the extent of MNase digestion (Figure S8B). Although the heterologous system used here may differ from endogenous chromatin assembly factors in *Oxytricha*, our experiment illustrates the principle that *trans*-acting factors can counteract or even overcome the effect of 6mA on nucleosome organization.

### Loss of 6mA impacts gene expression and sexual development

Recently, 6mA has been implicated in gene regulation in the green algae *Chlamydomonas* and basal yeasts because of its localization to TSSs of active genes (Fu et al., 2015; Mondo et al., 2017), similar to ciliates. However, this relationship has not been functionally tested. Since *mta1* mutants exhibit genome-wide loss of 6mA, we assayed these cells for transcriptional changes by poly(A)+ RNAseq. Only a small minority of genes show significant changes in gene expression (10% FDR, Figure 7A). To examine the methylation status of these differentially expressed genes, we grouped them according to “starting” methylation level, as defined by the total number of 6mA marks near the TSS in wild type cells. Genes exhibit two distinct transcriptional responses: those with low starting levels of 6mA exhibit a small change in 6mA between wild type and mutant cells (Figure 3D) and tend to be significantly upregulated in mutant lines (p = 2.8 × 10^−9^, Fisher’s exact test; and Figure 7B). Surprisingly, genes with high starting 6mA are not enriched in differentially expressed genes (p > 0.1, Fisher’s exact test), even though they exhibit greater loss of 6mA in mutants. Steady state RNAseq levels are therefore largely robust to drastic changes in 6mA levels. Since most, but not all, 6mA is lost from *mta1* mutants (Figure 3C), it is also possible that residual DNA methylation across the genome sufficiently buffers genes from changes in transcription.

**Figure 7.**
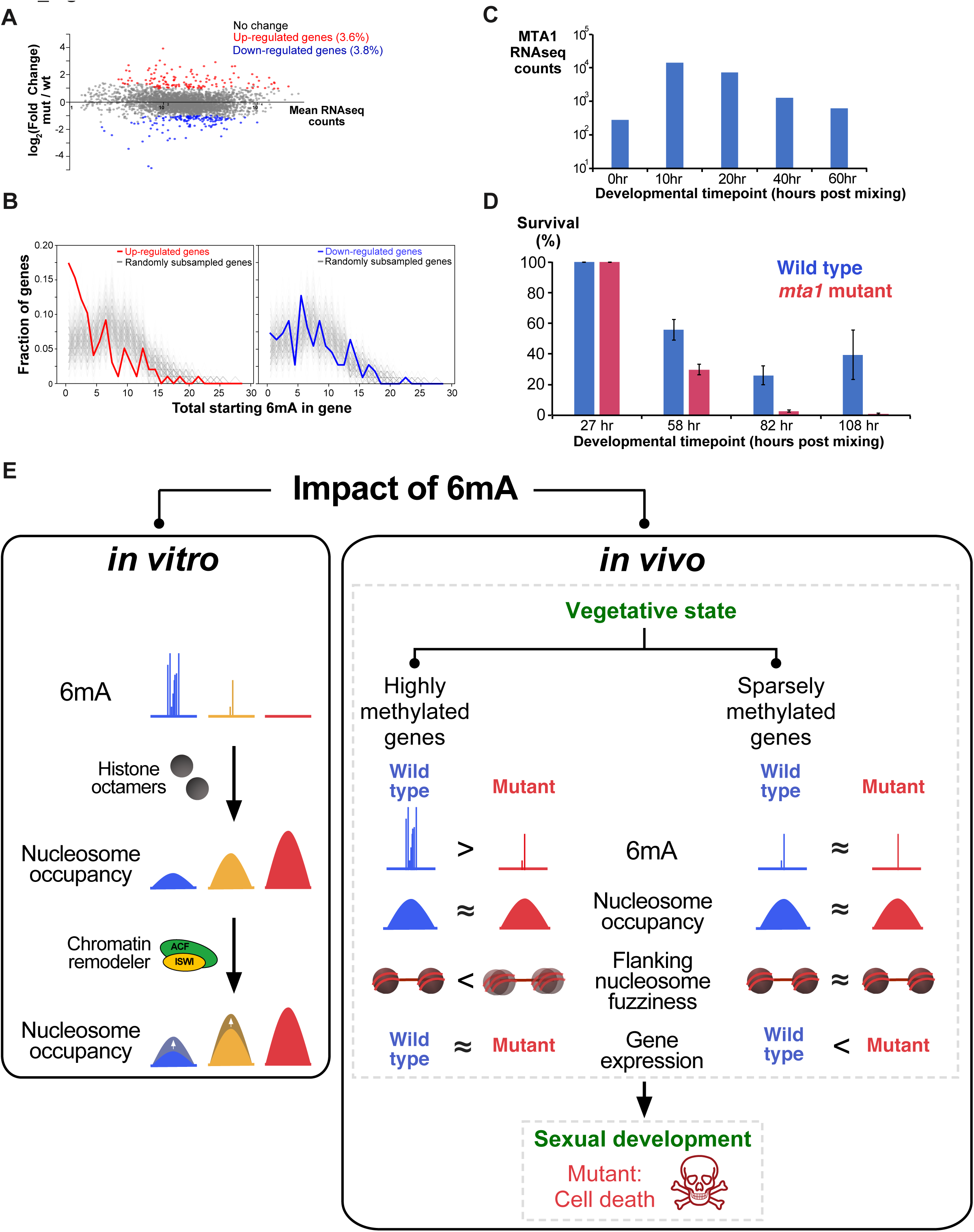
Effects of 6mA on gene expression and cell viability *in vivo*. A. RNAseq scatterplot showing a small percentage (<10%) of genes are differentially expressed in *mta1* mutants (top panel). Horizontal axis denotes the mean RNAseq counts across all biological replicates from wild type and mutant data for each gene. Vertical axis denotes the log_2_(fold change) in gene expression (mutant / wild type).
B. Upregulated genes tend to be sparsely methylated. The number of 6mA marks in upregulated and downregulated genes are plotted as red and blue lines, respectively. 6mA levels from an equal number of randomly subsampled, non-differentially regulated genes are plotted in grey. Subsampling is performed 1000 times, resulting in 1000 grey lines collectively visualized as a diffuse background pattern. 6mA levels between randomly sampled and differentially expressed genes are compared using a Wilcoxon rank-sum test. For upregulated genes, 894 out of 1000 significance tests showed p-value <= 0.05, while for downregulated genes, 0 out of 1000 significance tests showed p-value <= 0.05. 6mA levels in upregulated genes are therefore significantly different from that of control genes. *P*-values are corrected for multiple testing using the Benjamini-Hochberg method. Total starting 6mA level denotes the number of 6mA marks −100bp to +250bp from the TSS in wild type cells.
C. RNAseq analysis of MTA1 expression at various developmental timepoints during the sexual cycle of *Oxytricha*. RNAseq timecourse data are obtained from (Swart et al., 2013). *Oxytricha* cells from two different strains are mixed at 0 hr to induce mating and commence the sexual cycle. The total duration of the sexual cycle is ~60 hr.
D. Survival curve of *Oxytricha* cells during the sexual cycle. Cells are mixed at 0 hr to induce mating. Since not all cells enter the sexual cycle, mated cells are separated from unmated vegetative cells at 15 hr and transferred into a separate dish. The cells are allowed to rest for 12 hr to account for cell death during transfer. The number of surviving mated cells is counted from 27 hr onwards. The total cell number at each timepoint is normalized to 27 hr data to obtain the percentage survival. An increase in survival at 108 hr is observed in wild type samples because the cells have completed mating and reverted to the vegetative state, where they can proliferate and increase in number. Error bars in all panels represent s.e.m. (n = 4).
E. Summary model illustrating the impact of 6mA on nucleosome organization and gene expression. 6mA directly disfavors nucleosome occupancy *in vitro* in a quantitative, local manner. This effect can be overcome by the ATP-dependent action of chromatin remodelers such as ACF. In agreement with this observation *in vitro*, 6mA has minimal effect on nucleosome occupancy *in vivo*, where analogous protein complexes act on chromatin. Genes exhibit distinct responses to 6mA *in vivo*, depending on their starting methylation level. Highly methylated genes exhibit increases in nucleosome fuzziness upon 6mA loss, but do not change significantly in expression. In contrast, sparsely methylated genes are upregulated but do not show changes in nucleosome fuzziness. Loss of 6mA also leads to cell death during sexual development.

Because the aforementioned phenotypic changes were assayed in vegetative *Oxytricha* cells, we asked whether MTA1 may play roles outside of this developmental state. MTA1 transcript levels are markedly upregulated in the sexual cycle, as assayed by poly(A)+ RNAseq (Figure 7C). Strikingly, *mta1* mutants fail to complete the sexual cycle when induced to mate, and display complete lethality (Figure 7D). Thus, 6mA may function in a developmental stage-specific manner. Massive genome-wide DNA rearrangements occur during the *Oxytricha* sexual cycle, coincident with the upregulation of MTA1 (Bracht et al., 2013; Chen et al., 2014). It is possible that 6mA provides a bookmarking role in vegetative cells for subsequent global remodeling events. Our data warrant further investigation into the roles of 6mA in complex developmental processes.

## Discussion

Since the initial reports of 6mA in *C. elegans*, *Drosophila*, and *Chlamydomonas* in 2015, numerous studies have the documented the presence and genomic distribution of 6mA in diverse eukaryotic genomes. Two key questions remain: 1) What is the underlying machinery that reads, writes, and erases 6mA? 2) What is the function of 6mA? In this study, we used ciliates as a model system to address both problems. We first undertook an unbiased biochemical fractionation approach to discover putative 6mA methyltransferase(s). This led to the identification of MTA1 as a hitherto undescribed MT-A70 protein necessary for 6mA methylation in ciliates. It remains to be determined whether MTA1 is also necessary for m^6^A RNA methylation, and if so, what is the relationship between DNA and RNA modification. Remarkably, MTA1 does not belong to the METTL3 or METTL14 clades, which together constitute the canonical RNA m^6^A methyltransferase. It is also evolutionarily distinct from METTL4, a 6mA methyltransferase in *C. elegans* (Greer et al., 2015). Further biochemical and evolutionary studies are needed to shed light on its substrate specificity and any interacting partners that could modulate its activity.

We investigated the function of 6mA *in vitro* by building full-length, epigenetically defined chromosomes. This allowed us to show that 6mA directly disfavors nucleosome occupancy in a local, quantitative manner, independent of DNA sequence (Figure 7E). This effect can be overcome by ATP-dependent chromatin remodelers *in vitro*. We expect the biochemical impact of 6mA to be directly pertinent across a wide range of eukaryotic genomes, including vertebrates, *C. elegans*, *Drosophila* and fungi, where this epigenetic modification has recently been documented (Greer et al., 2015; Koziol et al., 2015; Liang et al., 2018; Liu et al., 2016; Mondo et al., 2017; Wu et al., 2016; Xiao et al., 2018; Yao et al., 2018; Zhang et al., 2015). Our experiments do not reveal exactly how 6mA disfavors nucleosome occupancy. Early studies suggest that 6mA destabilizes dA:dT base pairing, leading to a decrease in the melting temperature of DNA (Engel and von Hippel, 1978). Whether this or some other physico-chemical property of 6mA contributes to lowered nucleosome stability awaits further investigation.

To understand the biological roles of MTA1 *in vivo*, we analyzed nucleosome organization and gene expression in vegetative *mta1* mutant cells (Figure 7E). We find that nucleosome organization exhibits only subtle changes after genome-wide loss of 6mA. In addition, only a small set of genes (<10%) is transcriptionally dysregulated. It is possible that residual 6mA in *mta1* mutants could mask relevant phenotypes. Nonetheless, our results caution against interpreting 6mA function solely based on correlation with genomic elements. Recent studies have suggested that 6mA influences chromatin organization *in vivo*, based on the inverse relationship between 6mA and nucleosome positioning (Fu et al., 2015; Wang et al., 2017). Analogous observations in green algae show that cytosine methylation is clustered within nucleosome linkers, and that subsequent PCR amplification restores *in vitro* nucleosome occupancy (Huff and Zilberman, 2014). Our experiments indicate that 6mA also intrinsically disfavors nucleosomes *in vitro*, but – crucially – this effect can be overridden by distinct factors *in vitro* and *in vivo.* We propose that phased nucleosome arrays are first established *in vivo*, which then restrict MTA1-mediated methylation to linker regions due to steric hindrance. This in turn decreases the fuzziness of flanking nucleosomes, reinforcing chromatin organization. Therefore, 6mA tunes nucleosome organization *in vivo*. Our data do not support the hypothesis that nucleosome phasing is established by pre-deposited 6mA.

More broadly, we demonstrate that ciliate genomes – naturally abundant in 6mA – constitute a rich platform for uncovering novel enzymatic activities related to this epigenetic modification. Further mining of ciliate genomes will likely reveal unexpected biological functions of 6mA and the underlying machinery that guides these processes. Our work also showcases the utility of *Oxytricha* chromosomes for advancing chromatin biology. We demonstrate how key epigenetic features can be reconstructed using synthetic chromosomes with fully native DNA sequence. By extending current technologies (Müller et al., 2016), it should be feasible to introduce both modified nucleosomes and DNA methylation in a site-specific manner on full-length chromosomes. Such ‘designer’ chromosomes will serve as powerful tools for studying DNA-templated processes such as transcription within the context of a fully native DNA environment.

## Author contributions

L.Y.B. conceived the project, synthesized chromosomes, performed computational and experimental analysis for all Figures and Tables, and wrote the manuscript. G.T.D. synthesized chromosomes and prepared *Xenopus* histones. D.M.C. generated MTA1 mutants. R.E.T. synthesized 6mA nucleoside standards for mass spectrometry. K.A.L. processed raw SMRT-seq data. E.R.H. performed 6mA IP-seq. J.R.B. prepared *Oxytricha* DNA for SMRT-seq. R.P.S. performed SMRT-seq. T.W.M. and L.F.L. conceived the project and analyzed data. G.T.D, T.W.M, and L.F.L. edited the manuscript.

## Acknowledgments

We thank Geoffrey Dann for discussions regarding the ACF remodeler and for performing preliminary tests of its activity; Tharan Srikumar and Saw Kyin for assistance with nucleoside mass spectrometry; Istvan Pelczer for assistance with NMR measurements of nucleoside standards; Wei Wang, Jessica Wiggins, and Jennifer Miller for assistance with Illumina sequencing; C. David Allis, Virginia Zakian, Tim Bestor, and Samuel Sternberg for helpful discussions; Israel Fernandez and Alexander Sobolevsky for discussions and access to FPLC equipment; Krupa Jani for discussions regarding methyltransferase assays; Ana Mostafavi and Glen Liszczak for advice regarding amber codon suppression, and Jingmei Wang, Barbara Dul, and Fei Song for laboratory support. This work was funded by NIH grants R01-GM59708, R35-GM122555, and R01-GM109459 to L.F.L and R01-GM107047 to T.W.M.

---

**Table S1. Descriptive statistics**

**(A)** Properties of 6mA distribution in nucleosome linkers. In *Oxytricha*, methyl cluster 1 = between 5’ chromosome end and +1 nucleosome; methyl cluster 2 = between +1 and +2 nucleosome; methyl cluster 3 = between +2 and +3 nucleosome. In *Tetrahymena*, methyl cluster 1 = between +1 and +2 nucleosome; methyl cluster 2 = between +2 and +3 nucleosome; methyl cluster 3 = between +3 and +4 nucleosome. Consensus +1/+2/+3/+4 nucleosome positions: 193, 402, 618, 837 bp downstream of *Oxytricha* 5’ chromosome ends; 112, 304, 497, 698 bp downstream of *Tetrahymena* TSSs.

**(B)** Properties of *Oxytricha* chromosomes in native genomic DNA and mini-genome DNA. “+/-“ indicates one standard deviation above or below the mean.

**Table S2. Candidate 6mA methyltransferase genes in ciliates**

The Uniprot ID of each candidate gene is listed.

**Table S3. Protein sequences for phylogenetic tree construction**

FASTA-formatted list of amino acid sequences used to construct the phylogenetic tree in Figure 2A. Full headers and corresponding sequences were obtained from NCBI.

**Table S4. Primer sequences**

All primers are in the 5’ to 3’ direction.

**Figure S1. Mass spectrometry analysis confirms the presence of 6mA in ciliate DNA**

**(A)** *Oxytricha* and *Tetrahymena* genomic DNA were digested into nucleosides using degradase enzyme mix, followed by analysis using reverse-phase HPLC and mass spectrometry. Isotopically labeled dA and 6mA standards (^15^N_5_-dA and D_3_-6mA) were mixed with each sample to allow quantitative measurement of endogenous dA and 6mA concentrations. MS/MS analysis of labeled dA and 6mA standards confirmed the mass of the nucleobase. Eluted peaks with expected masses of dA and 6mA, and with highly similar retention times (RT) to internal standards are detected in *Oxytricha* and *Tetrahymena* nucleosides.

**(B)** Quantitation of dA and 6mA levels in *Oxytricha* and *Tetrahymena* gDNA using internal isotopically labeled nucleoside standards.

**Figure S2. Supplemental SMRT-seq data analyses**

**(A)** Top two panels depict PacBio coverage (horizontal axis) plotted against fractional methylation at each called 6mA site (vertical axis). Bottom left panel is a histogram of fractional methylation of all 6mA sites. Bottom right panel is a histogram of IPD ratios of all 6mA sites. Mutant datasets show significantly lower fractional methylation and IPD ratios at 6mA sites than wild type data.

**(B)** Wild type SMRT-seq data are randomly subsampled 15 times, such that the resulting coverage is lower than *mta1* mutant data. The difference in PacBio coverage between mutant and subsampled wildtype data is calculated for each chromosome, and is collectively represented as an olive boxplot (top panel). This set of calculations is repeated 15 times for each subsampled dataset, resulting in a series of 15 boxplots. The difference in PacBio coverage between mutant and fully sampled wild type data is represented as a violet boxplot. Separately, the difference in total 6mA marks per chromosome is calculated for respective datasets, and boxplots are shown in the bottom panel. Mutant datasets consistently yield lower numbers of called 6mA marks than subsampled wild type, despite the former having higher coverage than the latter.

**(C)** Scatterplot of total number of 6mA marks per chromosome in wild type versus mutant data. PacBio cutoffs for calling 6mA marks are varied as shown. A greater number of 6mA marks per chromosome are consistently detected in wild type than mutant data.

**(D)** Boxplot of PacBio chromosome coverage in individual wild type and mutant biological replicates (left panel). Only chromosomes with 100-150x PacBio coverage are shown. The total number of 6mA marks in each of these chromosomes are plotted in the right panel. Wild type replicates show consistently higher numbers of 6mA marks per chromosome than mutant replicates.

**Figure S3. Genomic distribution of 6mA in *Tetrahymena thermophila***

*Tetrahymena* MNase-seq data and RNA-seq data were obtained from previously published datasets (Beh et al., 2015; Xiong et al., 2012).

**(A)** Meta-chromosome plots overlaying *in vivo* MNase-seq (nucleosome occupancy) and SMRT-seq (6mA), relative to annotated transcription start sites. 6mA lies mainly within nucleosome linker regions, between the +1, +2, +3, and +4 nucleosomes.

**(B)** Histograms of the total number of 6mA marks within each linker in *Tetrahymena* genes. Calculations are performed as described in Figure 1B. Distinct linkers are highlighted with horizontal bold blue lines.

**(C)** Relationship between transcriptional activity and total number of 6mA marks in *Tetrahymena* genes. Analysis is performed as in Figure 1C.

**(D)** Composite analysis of 441,618 methylation sites reveals that 6mA occurs within an 5’-ApT-3’ dinucleotide motif in *Tetrahymena*, consistent with previous experiments (Bromberg et al., 1982; Wang et al., 2017) and similar to *Oxytricha*.

**(E)** Distribution of various 6mA dinucleotide motifs across the genome.

**(F)** Organization of transcription (mRNA-seq), nucleosome organization (MNase-seq), and 6mA (SMRT-seq) in a *Tetrahymena* gene.

**Figure S4. Domain organization and mass spectrometry coverage of MTA1**

**(A)** Domains in *Tetrahymena* and *Oxytricha* MTA1 proteins are predicted using hmmscan on the EMBL-EBI webserver, against the Pfam database (Finn et al., 2015). “aa” denotes amino acids. MT-A70 domains are colored in blue, while coiled coil motifs are colored in orange.

**(B)** LC-MS/MS coverage of MTA1. The full amino acid sequence of MTA1 is shown, and regions of the protein covered by peptide data are highlighted in red. “Low Salt Sample” and “High Salt Sample” correspond to partially purified nuclear extracts that elute as two distinct peaks of activity from a Q sepharose anion exchange column (Figure 2C). The Low Salt Sample and High Salt Sample are further purified through ion exchange and size exclusion chromatography to obtain fractions with low, medium, and high DNA methyltransferase activity (Figures 2C and 2G).

**Figure S5. Gene expression profile of *Tetrahymena* MTA1**

Microarray data are obtained from TetraFGD (Miao et al., 2009; Xiong et al., 2011). The horizontal axis categories beginning with “S” and “C” represent the number of hours since the onset of starvation and conjugation (mating), respectively. “Low”, “Med”, and “High” denote relative cell densities during log-phase growth. Blue and orange traces represent data from two biological replicates.

**Figure S6. Confirmation of ectopic DNA insertion in *mta1* mutants**

Poly(A)+ RNAseq analysis of wild type and *mta1* mutants. “ATG” denotes start codon of MTA1 gene. A 62bp ectopic DNA insertion results in a frameshift mutation in the MTA1 coding region. Three wild type (WT^1^, WT^2^, WT^3^) and mutant (*mta1^1^*, *mta1^2^*, *mta1^3^*) biological replicates are analyzed. Short horizontal bars represent RNAseq reads, which are ~75 nt in length and mapped to the reference sequence. For a read to be successfully mapped, it must have no more than 2 mismatches relative to the reference sequence. Unmapped reads are discarded. Blue and red bars denote RNAseq reads that map to native and ectopic regions, respectively. RNAseq reads overlapping the ectopic region are detected in mutant but not wild type replicates. These reads span junctions between the ectopic and flanking coding regions, confirming the site of ectopic insertion.

**Figure S7. Effects of 6mA loss on nucleosome organization *in vivo***

Nucleosomes are grouped according to their “starting” 6mA level, defined as the total number of 6mA marks +/-200 bp from the nucleosome dyad in wild type cells (WT). The dyad is assigned to be the peak position of MNase-seq reads. Similarly, linkers are grouped according to their “starting” methylation level, defined as the total number of 6mA marks between two flanking nucleosome dyads (or between the 5’ chromosome end and the terminal nucleosome) in wild type cells. Loci with high starting 6mA have methylation greater than or equal to the 90^th^ percentile of starting 6mA levels, and show greater changes in methylation between mutant and wild type cells (Figure 3D). Those with low starting 6mA are in the lowest 10^th^ percentile. If 6mA impacts nucleosome organization *in vivo*, then loci with high starting 6mA should show a greater change in nucleosome organization. Possible effects are illustrated in panels A – C. Vertical green lines depict 6mA marks, while blue and red peaks denote nucleosome occupancy. The plots shown in panels A – C illustrate the idealized result if 6mA disfavors nucleosomes *in vivo*. Actual effects are shown in panels D – G. “Wild type” is abbreviated as WT. Analyses are restricted to the 5’ chromosome end.

**(A)** 6mA loss may result in an increase in nucleosome fuzziness (highlighted with bold red double-sided arrow). The effect should be greater for nucleosomes with high starting 6mA due to greater change in 6mA between mutant and wild type cells (“Change in nucleosome fuzziness” Box). Nucleosomes should, in turn, exhibit lower occupancy near the peak position, and higher occupancy in flanking regions (“Change in Nucleosome occupancy” Box; highlighted with red arrowheads and plotted +/-73bp from the dyad). Nucleosome fuzziness is calculated as the standard deviation of MNase-seq read locations +/-73bp from the dyad.

**(B)** 6mA loss from nucleosome linker regions may result in a decrease in linker length (highlighted with bold red bracket). If so, the magnitude of decrease in linker length should be greater for linkers with high starting 6mA (“Change in linker length” Box).

**(C)** 6mA loss may result in an increase in occupancy directly over the methylated linker region (highlighted with bold red bracket). If so, the magnitude of increase in linker occupancy should be greater for regions with high starting 6mA (“Change in linker occupancy” Box). Linker occupancy denotes the average MNase-seq coverage +/-25bp from the midpoint between flanking nucleosome dyads or chromosome end. As an example, for the +1/+2 nucleosome linker, occupancy is calculated +/-25bp from the midpoint of the +1 and +2 nucleosome dyad positions. Since nucleosome linker length in *Oxytricha* is ~200bp (Figure S9), the genomic window used to calculate linker occupancy has minimal overlap with that for calculating nucleosome fuzziness and occupancy in panel A.

**(D)** *Impact of 6mA loss on nucleosome fuzziness*. For each nucleosome, the change in fuzziness between mutant and wild type cells is calculated. Boxplots represent the distribution of changes in fuzziness scores. “MNase-seq” denotes sequencing of nucleosomal DNA obtained from *Oxytricha* chromatin *in vivo*, while “Control gDNA-seq” represents sequencing of MNase-digested, naked genomic DNA *in vitro*. Notch in the boxplot denotes median, ends of boxplot denote first and third quartiles, upper whisker denotes third quantile + 1.5 × interquartile range, and lower whisker denotes data quartile 1 − 1.5 × interquartile range. Distributions are compared using a Wilcoxon rank-sum test. N.S denotes “non-significant”, with p > 0.01.

**(E)** *Impact of 6mA loss on nucleosome occupancy*. For each nucleosome, the difference in nucleosome occupancy between mutant and wild type cells is calculated at individual basepairs +/-73bp around the nucleosome dyad. Data are averaged and depicted as line plots. The change in occupancy at the dyad is compared between nucleosomes with high and low starting 6mA using a Wilcoxon rank-sum test.

**(F)** *Impact of 6mA loss on linker length.* Three types of linkers are analyzed: between the 5’ chromosome end and +1 nucleosome dyad, between the +1 and +2 nucleosome dyads, and between the +2 and +3 nucleosome dyads. For each linker, the difference in its length between mutant and wild type cells is calculated. The resulting distribution of linker length differences is plotted as a histogram. Distributions are compared using two-sample unequal variance t-test. N.S. indicates “not significant”, with p > 0.01.

**(G)** *Impact of 6mA loss on linker occupancy.* Linkers are binned as in panel F. For each linker, the difference in occupancy between mutant and wild type cells is calculated. The resulting distribution of changes in linker occupancy is represented as a boxplot. Notch in the boxplot denotes median, ends of boxplot denote first and third quartiles, upper whisker denotes third quantile + 1.5 × interquartile range, and lower whisker denotes data quartile 1 − 1.5 × interquartile range. Distributions are compared using two-sample unequal variance t-test. N.S. indicates “not significant”, with p > 0.01.

**Figure S8. MNase-seq analysis is robust to variation in MNase digestion**

(A) Same analysis as Figure 4C, showing that 6mA quantitatively disfavors nucleosome occupancy *in vitro* but not *in vivo*. Here, the extent of MNase digestion was 40% of that in Figure 4C. *P*-values were calculated using a two-sample unequal variance t-test. N.S denotes “non-significant”, with p > 0.05.

(B) Same analysis as Figure 6E, showing that the ACF complex restores nucleosome occupancy over methylated DNA in an ATP-dependent manner *in vitro*. Here, the extent of MNase digestion was 25% of that in Figure 6E. *P*-values were calculated using a two-sample unequal variance t-test. N.S denotes “non-significant”, with p > 0.05.

(C) Same analysis as Figure S7D, showing that nucleosomes with high starting 6mA show larger changes in fuzziness. Here, the extent of MNase digestion was 40% of that in Figure S7D. Distributions are compared using a Wilcoxon rank-sum test. N.S denotes “non-significant”, with p > 0.01.

(D) Same analysis as Figure S7E, showing that nucleosomes with high starting 6mA exhibit characteristic changes in nucleosome occupancy at and around the nucleosome dyad. Here, the extent of MNase digestion was 40% of that in Figure S7E. The change in dyad occupancy is compared between nucleosomes with high and low starting 6mA using a Wilcoxon rank-sum test. N.S denotes “non-significant”, with p > 0.01.

**Figure S9. Distribution of nucleosome linker lengths *in vivo***

Linker types analyzed: between the 5’ chromosome end and +1 nucleosome dyad, between the +1 and +2 nucleosome dyads, and between the +2 and +3 nucleosome dyads. Analysis here is restricted to the 5’ chromosome end. Orange ovals represent heterodimeric telomere end-binding proteins that protect the chromosome end *in vivo*. Each linker type is further divided according to the level of starting 6mA (high/low) as defined in Figure S7. The length distribution of each group of linkers in wild type and mutant cells is depicted as overlaid histograms. The median linker length from each group is included as an inset. Distributions are compared using a Wilcoxon rank-sum test. N.S. indicates “not significant”, with p > 0.05.

**Figure S10. Preparation of histone octamers for chromatin assembly**

*Xenopus* unmodified core histones were recombinantly expressed. *Oxytricha* histones were acid-extracted from vegetative nuclei. *Oxytricha* and *Xenopus* histones were subsequently refolded into octamers and purified through size exclusion chromatography.

**(A)** Reverse-phase HPLC purification of acid-extracted *Oxytricha* histones. Fractions 1-5 were individually collected and analyzed by Coomassie staining and western blotting.

**(B)** SDS-PAGE analysis of purified *Oxytricha* histone fractions.

**(C)** Western blot analysis of *Oxytricha* histone fractions 1-5. The fraction that is most enriched in each type of histone is colored in red. Arrowheads indicate likely histone bands.

**(D)** SDS-PAGE analysis of purified *Oxytricha* and *Xenopus* histone octamers.

**Figure S11. Gel analysis of assembled chromatin**

*Xenopus* or *Oxytricha* histone octamers were assembled on DNA and subsequently digested with MNase to obtain ~150bp mononucleosome-sized fragments (labeled with red arrowheads). The resulting products were visualized by agarose gel electrophoresis. Mononucleosomal DNA was gel-excised and analyzed using Illumina sequencing or tiling qPCR analysis as described in Figures 4, 6, S8 and S13.

**(A)** Chromatin was assembled on PCR-amplified *Oxytricha* mini-genome DNA, digested with MNase, and analyzed by agarose gel electrophoresis.

**(B)** Chromatin was assembled on native *Oxytricha* genomic DNA, digested with MNase, and analyzed by agarose gel electrophoresis.

**(C)** Chromatin was assembled with synthetic chromosome DNA, digested with MNase, and visualized by agarose gel electrophoresis. All assemblies with synthetic chromosomes were performed in the presence of an approximately 100-fold mass excess of buffer DNA relative to synthetic chromosome (see Methods). This applies to panels C, D, and E. Representative assemblies with the unmethylated chromosome are shown. Methylated chromosome assemblies were separately performed in place of the unmethylated variant.

**(D)** Chromatin was assembled on unmethylated synthetic chromosomes by salt dialysis and subsequently incubated with ACF and/or ATP. The resulting mixture was digested with MNase and visualized by agarose gel electrophoresis. Regularly spaced nucleosomes (labeled with red dots) are observed only when chromatin was incubated with both ACF and ATP.

**(E)** Chromatin assembled on unmethylated synthetic chromosomes using the NAP1 histone chaperone in the presence of ACF and/or ATP. The resulting mixture was digested with MNase and visualized by agarose gel electrophoresis. Nucleosomes are regularly spaced (labeled with red dots) in the presence of both ACF and ATP, although less apparent than in panel D.

**Figure S12. Detailed chromosome synthesis scheme**

Staggered dotted lines represent *BsaI* cleavage sites. Chromosomes are numbered according to Figures 5B and 5C.

**(A)** Assembly of synthetic chromosomes bearing 6mA at cognate sites. PCR-amplified building blocks were digested with *BsaI* and subsequently purified. They were then mixed with a short linker of annealed oligonucleotides and ligated to form the full-length chromosome.

**(B)** Assembly of synthetic chromosomes bearing 6mA at ectopic sites. Individual PCR-amplified building blocks were digested with *BsaI* and subsequently purified. *BsaI* digestion and *EcoGII* methylation were performed in the same reaction. All building blocks were subsequently ligated to obtain the full-length chromosome.

**(C)** Sanger sequencing of cloned synthetic chromosomes. Continuous horizontal green bars represent full sequence identity between the reference chromosome and individual sequencing reads. All synthetic chromosomes show the expected sequence with correct ligation junctions.

**Figure S13. Supplemental qPCR analysis**

**(A)** Tiling qPCR analysis of nucleosome occupancy in spike-in and homogeneous synthetic chromosome preparations. The blunt, unmethylated synthetic chromosome (construct #1 in Figure 5B) was used for chromatin assembly with (“Spike-in”) or without (“Homogeneous”) a 100-fold excess of buffer DNA. In the latter case, an equivalent mass of synthetic chromosome was added in place of buffer DNA to maintain the same DNA concentration for chromatin assembly. The tiling qPCR assay was performed as in Figure 6B. Shaded red bars depict the regions where 6mA modulates nucleosome occupancy in separate methylated chromosomes analyzed in Figures 6B and 6C. Note that methylated chromosomes were not used to generate qPCR data for this Figure. Black arrowheads indicate no decrease in nucleosome occupancy in these regions when buffer DNA is used. Thus, the decrease in nucleosome occupancy in methylated chromosomes reported in Figure 6 cannot be attributed to spike-in versus homogeneous addition of DNA for chromatin assembly. Error bars in all panels represent s.e.m. (n = 3-4).

**(B)** Chromatin was assembled on synthetic chromosomes using the NAP1 histone chaperone in the presence of ACF and/or ATP, instead of salt dialysis. qPCR analysis was performed as in Figure 6B. Methylated chromosomes used in this experiment contain 6mA in native sites. The addition of ACF and ATP results in a partial restoration of nucleosome occupancy over the methylated region. These results are similar to Figure 6D, where chromatin was assembled by salt dialysis instead of NAP1.

**Figure S14. Putative ciliate ISWI orthologs**

ISWI is a member of the SW12/SNF2 ATPase family that acts as chromatin remodelers. The *Oxytricha* and *Tetrahymena* genomes were queried by BLASTP using *Drosophila melanogaster* ISWI (UniProt ID: Q24368), and the reciprocal best hit was retrieved from each genome. BLASTP E-values ≈ 0 in each case. Putative *Tetrahymena* and *Oxytricha* orthologs were queried for protein domains and associated GO terms using InterPro (Finn et al., 2017). Positions of predicted domains are depicted as horizontal orange bars on the left of the Figure, together with corresponding domain names and InterPro IDs. GO terms associated with the protein are listed on the right. ISWI contains an N-terminal catalytic ATPase domain and a C-terminal HAND-SANT-SLIDE module necessary for nucleosome binding and mobilization.

## STAR Methods

### Contact for reagent and resource sharing

Further information and requests for resources and reagents should be directed to and will be fulfilled by the Lead Contact, Laura Landweber.

### Experimental model and subject details

#### Oxytricha trifallax

Vegetative *Oxytricha trifallax* strain JRB310 was cultured at a density of 1.5 × 10^7^ cells/L to 2.5 × 10^7^ cells/L in Pringsheim media (0.11mM Na_2_HPO_4_, 0.08mM MgSO_4_, 0.85mM Ca(NO_3_)_2_, 0.35mM KCl, pH 7.0) and fed daily with *Chlamydomonas reinhardtii.* Cells were filtered through cheesecloth to remove debris and collected on a 10μm Nitex mesh for subsequent experiments.

#### Tetrahymena thermophila

Vegetative *Tetrahymena thermophila* strain SB210 were cultured as previously described (Beh et al., 2015). Stock cultures were maintained in Neff medium (0.25% w/v proteose peptone, 0.25% w/v yeast extract, 0.5% glucose, 33.3µM FeCl_3_) or propagated to high density in SSP medium (2% w/v proteose peptone, 0.1% w/v yeast extract, 0.2% w/v glucose, 33µM FeCl_3_) for nuclear extract preparation. Cells were collected by centrifugation for subsequent experiments.

### Method details

#### in vivo MNase-seq

3 × 10^5^ vegetative *Oxytricha* cells were fixed in 1% (w/v) formaldehyde for 10 min at room temperature with gentle shaking, and then quenched with 125mM glycine. Macronuclei were subsequently isolated by sucrose gradient centrifugation from fixed cells as previously described (Lauth et al., 1976). Purified nuclei were pelleted by centrifugation at 4000 × g, washed in 50ml TMS buffer (10mM Tris pH 7.5, 10mM MgCl_2_, 3mM CaCl_2_, 0.25M sucrose), resuspended in a final volume of 300μL, and equilibriated at 37°C for 5 min. Chromatin was then digested with MNase (New England Biolabs) at a final concentration of 15.7 Kunitz Units / μL at 37°C for 1 min 15 sec, 3 min, 5 min, 7 min 30sec, 10 min 30 sec, and 15 min respectively. Reactions were stopped by adding 1/2 volume of PK buffer (300mM NaCl, 30mM Tris pH 8, 75mM EDTA pH 8, 1.5% (w/v) SDS, 0.5mg/mL Proteinase K). Each sample was incubated at 65°C overnight to reverse crosslinks and deproteinate samples. Subsequently, nucleosomal DNA was purified through phenol:chloroform:isoamyl alcohol extraction and ethanol precipitation. Each sample was loaded on a 2% agarose-TAE gel to check the extent of MNase digestion. The sample exhibiting ~80% mononucleosomal species was selected for MNase-seq analysis, in accordance with previous guidelines (Zhang and Pugh, 2011). Mononucleosome-sized DNA was gel-purified using a QIAquick gel extraction kit (QIAGEN). Illumina libraries were prepared using an NEBNext Ultra II DNA Library Prep Kit (New England Biolabs) and subjected to paired-end sequencing on an Illumina HiSeq 2500 according to manufacturer’s instructions. All *Tetrahymena* MNase-seq data were obtained from (Beh et al., 2015).

#### poly(A)+ RNA-seq and 5’-complete cDNA-seq

*Oxytricha* cells were lysed in TRIzol reagent (Thermo Fisher Scientific) for total RNA isolation according to manufacturer’s instructions. Poly(A)+ RNA was then purified using the NEBNext Poly(A) mRNA Magnetic Isolation Module (New England Biolabs). The *Oxytricha* poly(A)+ RNA was prepared for RNA-seq using the ScriptSeq v2 RNA-Seq Library Preparation Kit (Illumina). *Tetrahymena* poly(A)+ RNA-seq data was obtained from (Xiong et al., 2012). The 5’ ends of capped RNAs were enriched from vegetative *Oxytricha* total RNA using the RAMPAGE protocol (Batut et al., 2013), and used for library preparation, Illumina sequencing and subsequent transcription start site determination (ie. “TSS-seq”). *Tetrahymena* transcription start site positions were obtained from (Beh et al., 2015). TSS positions used in analysis outside of Figure 1A were obtained from (Swart et al., 2013) and (Beh et al., 2015).

For RNAseq analysis of genes grouped according to “starting” methylation level level: total 6mA was counted between 100 bp upstream to 250 bp downstream of the TSS. Genes with high starting methylation have total 6mA in the 90^th^ percentile and higher. Genes with low starting methylation have total 6mA at or below the 10^th^ percentile.

#### 6mA IP-seq

Genomic DNA was isolated from vegetative *Oxytricha* cells using the Nucleospin Tissue Kit (Takara Bio USA, Inc.). DNA was sheared into 150bp fragments using a Covaris LE220 ultra-sonicator (Covaris). Samples were gel-purified on a 2% agarose-TAE gel, blunted with DNA polymerase I (New England Biolabs), and purified using MinElute spin columns (QIAGEN). The fragmented DNA was dA-tailed using Klenow Fragment (3’ -> 5’ exo-) (New England Biolabs) and ligated to Illumina adaptors following manufacturer’s instructions. Subsequently, 2.2μg of adaptor-ligated DNA containing 6mA was immunoprecipitated using an anti-N6-methyladenosine antibody (Cedarlane Labs) conjugated to Dynabeads Protein A (Invitrogen). The anti-6mA antibody is commonly used for RNA applications, but has also been demonstrated to recognize 6mA in DNA (Fioravanti et al., 2013; Xiao and Moore, 2011). The immunoprecipitated and input libraries were treated with proteinase K, extracted with phenol:chloroform, and ethanol precipitated. Finally, they were PCR-amplified using Phusion Hot Start polymerase (New England Biolabs) and used for Illumina sequencing.

#### Sample preparation for SMRT-seq

Macronuclei were isolated from vegetative *Oxytricha* and *Tetrahymena* as previously described (Lauth et al., 1976) and used for genomic DNA isolation with the Nucleospin Tissue Kit (Macherey-Nagel). Alternatively, whole *Oxytricha* cells instead of macronuclei were used. SMRT-seq was performed as previously described (Chen et al., 2014), according to manufacturer’s instructions, using P5-C3 and P6-C4 chemistry. *Oxytricha* and *Tetrahymena* macronuclear DNA were used for SMRT-seq in Figure 1 and S3, while *Oxytricha* whole cell DNA was used for all other Figures. Since almost all DNA in *Oxytricha* cells is derived from the macronucleus (Prescott, 1994), similar results are expected between the use of purified macronuclei or whole cells.

#### Illumina data processing

Reads from all biological replicates were merged before downstream processing. All Illumina sequencing data were quality trimmed (minimum quality score = 20) and length-filtered (minimum read length = 40nt) using Galaxy (Blankenberg et al., 2010; Giardine et al., 2005; Goecks et al., 2010). MNase-seq and 6mA IP-seq reads were mapped to complete chromosomes in the *Oxytricha trifallax* JRB310 (August 2013 build) or *Tetrahymena* thermophila SB210 macronuclear reference genomes (June 2014 build) using Bowtie2 (Langmead and Salzberg, 2012) with default settings, while poly(A)+ RNA-seq and TSS-seq reads were mapped using TopHat2 (Mortazavi et al., 2008) with August 2013 *Oxytricha* gene models or June 2014 *Tetrahymena* gene models, with default settings.

Within each MNase-seq dataset, the read pair length of highest frequency was identified. All read pairs with length +/-25bp from this maximum were used for downstream analysis. 6mA IP-seq single-end reads were extended to the mean fragment size, computed using cross-correlation analysis (Kharchenko et al., 2008). The per-basepair coverage of *Oxytricha* MNase-seq read pair centers (termed “nucleosome occupancy” in this manuscript) and extended 6mA IP-seq reads were respectively computed across the genome. Subsequently, the per-basepair coverage values were normalized by the average coverage within each chromosome to account for differences in DNA copy number (and hence, read depth) between *Oxytricha* chromosomes (Swart et al., 2013). The per-basepair coverage values were then smoothed using a Gaussian filter of standard deviation = 15. For RNA-seq data, the number of reads per kilobase of chromosome per million mapped reads (RPKM) was calculated for each chromosome without normalization by DNA copy number since there is no correlation between *Oxytricha* DNA and transcript levels (Swart et al., 2013). *Oxytricha* TSS-seq data were processed using CAGEr (Haberle et al., 2015); with clusterCTSS parameters (threshold = 1.6, thresholdIsTpm = TRUE, nrPassThreshold = 1, method = “paraclu”, removeSingletons = TRUE, keepSingletonsAbove = 5). Only TSSs with tags per million counts > 0.1 were used for downstream analysis. *Tetrahymena* MNase-seq data were processed similarly to *Oxytricha*, except that DNA copy number normalization was omitted as *Tetrahymena* chromosomes have uniform copy number (Eisen et al., 2006).

Nucleosome dyads were called as local maxima in MNase-seq coverage, as previously described (Beh et al., 2015). “Consensus” +1, +2, +3 nucleosome positions downstream of the TSS were inferred from aggregate MNase-seq profiles across the genome (Figure 1A for *Oxytricha* and Figure S3 for *Tetrahymena*). Each gene was classified as having a +1, +2, +3 and/or +4 nucleosome if there is a called nucleosome dyad within 75 bp of the consensus nucleosome position.

#### SMRT-seq data processing

We processed SMRT-seq data with SMRTPipe v1.87.139483 in the SMRT Analysis 2.3.0 environment using, in order, the P_Fetch, P_Filter (with minLength = 50, minSubreadLength = 50, readScore = 0.75, and artifact = −1000), P_FilterReports, P_Mapping (with gff2Bed = True, pulsemetrics = DeletionQV, IPD, InsertionQV, PulseWidth, QualityValue, MergeQV, SubstitutionQV, DeletionTag, and loadPulseOpts = byread), P_MappingReports, P_GenomicConsensus (with algorithm = quiver, outputConsensus = True, and enableMapQVFilter = True), P_ConsensusReports, and P_ModificationDetection (with identifyModifcations = True, enableMapQVFilter = False, and mapQvThreshold = 10) modules. All other parameters were set to the default. The *Oxytricha* August 2013 reference genome build was used for mapping *Oxytricha* SMRT-seq reads, with Contig10040.0.1, Contig1527.0.1, Contig4330.0.1, and Contig54.0.1 removed, as they are perfect duplicates of other Contigs in the assembly. *Tetrahymena* SMRT-seq reads were mapped to the June 2014 reference genome build. Only chromosomes with high SMRT-seq coverage (>= 80x for *Oxytricha*; >=100x for *Tetrahymena*) were used for all 6mA-related analyses.

#### Chromosome synthesis

Synthetic Contig1781.0 chromosomes were constructed from “building blocks” of native chromosome sequence (Figures 5B, 5C, and S12). These chromosome segments were generated from either annealed synthetic oligonucleotides or from genomic DNA via PCR-amplification using Phusion DNA polymerase (New England Biolabs). The latter contains terminal restriction sites for *BsaI* (New England Biolabs), a type IIS restriction enzyme that cuts distal from these sites. *BsaI* cleaves within the native DNA sequence, generating custom 4nt 5’ overhangs and releasing the non-native *BsaI* restriction site as small fragments that are subsequently purified away. The *BsaI*-generated overhangs are complementary only between adjacent building blocks, conferring specificity in ligation and minimizing undesired by-products. After *BsaI* digestion, PCR building blocks were purified by phenol:chloroform extraction and ethanol precipitation. Building blocks were then sequentially ligated to each other as described in Figure S12 using T4 DNA ligase (New England Biolabs) and purified by phenol:chloroform extraction and ethanol precipitation. Size selection after each ligation step was performed using polyethylene glycol (PEG) precipitation or Ampure XP beads (Beckman Coulter) to enrich for the large ligated product over its smaller constituents. The size of individual building blocks and their corresponding order of ligation were designed to maximize differences in size between ligated products and individual building blocks. This increases the efficiency in size selection of products over reactants. 6mA was installed in synthetic chromosomes using annealed oligonucleotides, or by incubation of DNA building blocks with *EcoGII* methyltransferase (New England Biolabs).

#### Verification of synthetic chromosome sequences

All chromosomes were dA-tailed using Klenow Fragment (3’ -> 5’ exo-) (New England Biolabs), cloned using a TOPO TA cloning kit (Thermo Fisher) or StrataClone PCR Cloning Kit (Agilent Technologies), transformed into One Shot TOP10 chemically competent *E. coli*, and sequenced using flanking T7, T3, M13F, or M13R primers.

#### Preparation of *Oxytricha* histones

Vegetative *Oxytricha trifallax* strain JRB310 was cultured as previously described (Swart et al., 2013). Cells were starved for 14 hr and subsequently harvested for macronuclear isolation as previously described (Lauth et al., 1976). Purified nuclei were pelleted by centrifugation at 4000 × g, resuspended in 0.421mL 0.4N H_2_SO_4_ per 10^6^ input cells, and nutated for 3 hr at 4°C to extract histones. Subsequently, the acid-extracted mixture was centrifuged at 21,000 × g for 15 min to remove debris. Proteins were precipitated from the cleared supernatant using trichloroacetic acid (TCA), washed with cold acetone, then dried and resuspended in 2.5% (v/v) acetic acid. Individual core histone fractions were purified from crude acid-extracts using semi-preparative RP-HPLC (Vydac C18, 12 micron, 10 mM × 250 mm) with 40-65% HPLC solvent B over 50 min (Figure S10A). The identity of each purified histone fraction was verified by western blotting (Figure S10C) using antibodies: anti-H2A (Active Motif #39111), anti-H2B (Abcam #ab1790), anti-H3 (Abcam #ab1791), anti-H4 (Active Motif #39269).

#### Preparation of recombinant *Xenopus* histones

All RP-HPLC analyses were performed using 0.1% TFA in water (HPLC solvent A), and 90% acetonitrile, 0.1% TFA in water (HPLC solvent B) as the mobile phases. Wild-type *Xenopus* H4, H3 C110A, H2B and H2A proteins were expressed in BL21(DE3) pLysS *E.coli* and purified as previously described (Debelouchina et al., 2016). Purified histones were characterized by ESI-MS using a MicrOTOF-Q II ESI-Qq-TOF mass spectrometer (Bruker Daltonics). H4: calculated 11,236 Da, observed 11,236.1 Da; H3 C110A: calculated 15,239 Da, observed 15,238.7 Da; H2A: calculated 13,950 Da, observed 13,949.8 Da; H2B: calculated 13,817 Da, observed 13,816.8 Da.

#### Preparation of histone octamers

*Oxytricha* and *Xenopus* histone octamers were respectively refolded from core histones using established protocols (Beh et al., 2015; Debelouchina et al., 2016). Briefly, lyophilized histone proteins (*Xenopus* modified or wild-type; *Oxytricha* acid-extracted) were combined in equimolar amounts in 6 M guanidine hydrochloride, 20 mM Tris pH 7.5 and the final concentration was adjusted to 1mg/mL. The solution was dialyzed against 2M NaCl, 10mM Tris, 1mM EDTA, and the octamers were purified from tetramer and dimer species using size-exclusion chromatography on a Superdex 200 10/300 column (GE Healthcare Life Sciences). The purity of each fraction was analyzed by SDS-PAGE. Pure fractions were combined, concentrated and stored in 50% v/v glycerol at −20°C.

#### Preparation of mini-genome DNA

98 full-length chromosomes were individually amplified from *Oxytricha trifallax* strain JRB310 genomic DNA using Phusion DNA polymerase (New England Biolabs). Primer pairs are listed in Table S4. Amplified chromosomes were separately purified using a MinElute PCR purification kit (QIAGEN), and then mixed in equimolar ratios to obtain “mini-genome” DNA. The sample was concentrated by ethanol precipitation and adjusted to a final concentration of ~1.6mg/mL.

#### Preparation of native genomic DNA for chromatin assembly

Macronuclei were isolated from vegetative *Oxytricha trifallax* strain JRB310 as previously described (Lauth et al., 1976), and genomic DNA was isolated using the Nucleospin Tissue kit (Macherey-Nagel). Approximately 200µg of genomic DNA was loaded on a 15%-40% linear sucrose gradient and centrifuged in a SW 40 Ti rotor (Beckman Coulter) at 160,070 × g for 22.5hr at 20°C. Sucrose solutions were in 1M NaCl, 20mM Tris pH 7.5, 5mM EDTA. Individual fractions from the sucrose gradient were analyzed on 0.9% agarose-TAE gels. Fractions containing high molecular weight DNA that migrated at the mobility limit were discarded as such DNA species were found to interfere with downstream chromatin assembly. All other fractions were pooled, ethanol precipitated, and adjusted to 0.5mg/mL DNA.

#### Chromatin assembly and preparation of mononucleosomal DNA

Chromatin assemblies were prepared by salt gradient dialysis as previously described (Beh et al., 2015; Luger et al., 1999), or using the NAP1 histone chaperone and ACF chromatin remodeler as previously described (An and Roeder, 2003; Fyodorov and Kadonaga, 2003). To reduce sample requirements while maintaining adequate DNA concentrations for chromatin assembly, synthetic chromosomes were first mixed with a hundred-fold excess of “buffer” DNA (PCR-amplified *Oxytricha* Contig17535.0). We verified that nucleosome occupancy in terminal regions (qPCR primer pairs 1 and 24) and the methylated region (qPCR primer pairs 6 and 7) of the synthetic chromosome is unaffected by the presence of buffer DNA (Figure S13A). Native and mini-genome DNA were not mixed with buffer DNA prior to chromatin assembly.

For chromatin assembly through salt dialysis: histone octamers and (synthetic chromosome + buffer) DNA were mixed in a 0.8:1 mass ratio, while histone octamers and (native or mini-genome) DNA were mixed in a 1.3:1 mass ratio, each in a 50μL total volume. Samples were first dialyzed into start buffer (10mM Tris pH 7.5, 1.4M KCl, 0.1mM EDTA pH 7.5, 1mM DTT) for 1 hr at 4°C. Then, 350mL end buffer (10mM Tris pH 7.5, 10mM KCl, 0.1mM EDTA, 1mM DTT) was added at a rate of 1mL/min with stirring. The assembled chromatin was dialyzed overnight at 4°C into 200mL end buffer, followed by a final round of dialysis in fresh 200mL end buffer for 1 hr at 4 °C. The assembled chromatin was then adjusted to 50mM Tris pH 7.9, 5mM CaCl_2_ and digested with MNase (New England Biolabs) to mainly mononucleosomal DNA as previously described (Beh et al., 2015).

For chromatin assembly using NAP1 and ACF: 0.49µM NAP1 and 58nM histone octamer were first mixed in a 302µl reaction volume containing 62mM KCl, 1.2% (w/v) polyvinyl alcohol (Sigma Aldrich), 1.2% (w/v) polyethylene glycol 8000 (Sigma Aldrich), 25mM HEPES-KOH pH 7.5, 0.1mM EDTA-KOH, 10% (v/v) glycerol, and 0.01% (v/v) NP-40. The NAP1-histone mix was incubated on ice for 30 min. Meanwhile, “AM” mix was prepared, consisting of 20mM ATP (Sigma Aldrich), 200mM creatine phosphate (Sigma Aldrich), 33.3mM MgCl_2_, 33.3µg/µl creatine kinase (Sigma Aldrich) in a 56µl reaction volume. After the 30 min incubation, 5.29 µl of 1.7 µM ACF complex (Active Motif) and the “AM” mix were sequentially added to the NAP1-histone mix. Then, 10.63µl of native or mini-genome DNA (2.66µg) was added, resulting in a 374µl reaction volume. The final mixture was incubated at 27°C for 2.5 hr to allow for chromatin assembly. Subsequently, CaCl_2_ was added to a final concentration of 5mM, and the chromatin was digested with MNase (New England Biolabs) to mainly mononucleosomal DNA as previously described (Beh et al., 2015).

Mononucleosome-sized DNA from MNase-digested chromatin was gel-purified and used for tiling qPCR on a Viia 7 Real-Time PCR System with Power SYBR Green PCR master mix (Thermo Fisher), or *in vitro* MNase-seq on an Illumina HiSeq 2500, according to the manufacturer’s instructions. qPCR primer sequences are listed in Table S4.

#### Tiling qPCR analysis

qPCR data were analyzed using the ΔΔCt method (Livak and Schmittgen, 2001). At each locus along the synthetic chromosome, ΔCt = (Ct at locus of interest) – (Ct at qPCR primer pair 22, far from the methylated region). See Figure 6B for location of qPCR primer pair 22. Separate ΔCt values were calculated from mononucleosomal DNA and the corresponding naked, undigested synthetic chromosome. The ΔΔCt value was calculated from this pair of ΔCt values. This controls for potential variation in PCR amplification efficiency, especially over methylated regions. The fold change in mononucleosomal DNA relative to naked chromosomal DNA at a particular locus is calculated as 2^-ΔΔCt^, and denotes ‘nucleosome occupancy’ for all presented qPCR data.

#### ACF spacing assay

ATP-dependent nucleosome spacing was performed in accordance with a previous study (Lieleg et al., 2015). Chromatin was assembled by salt gradient dialysis as described above, and then adjusted to 20mM HEPES-KOH pH 7.5, 80mM KCl, 0.5mM EGTA, 12% (v/v) glycerol, 10mM (NH_4_)_2_SO_4_, 2.5mM DTT. Samples were then incubated for 2.5 hr at 27°C with 3mM ATP, 30mM creatine phosphate, 4mM MgCl_2_, 5 ng/μl creatine kinase, and 11 ng/μL ACF complex (Active Motif). Remodeled chromatin was then adjusted to 5mM CaCl_2_ and subjected to MNase digestion, mononucleosomal DNA purification, and qPCR analysis as described above.

#### Phylogenetic analysis of MT-A70 proteins

The MTA1 amino acid sequence (XP_001032074.3) was queried against the NCBI nr database using PSI-BLAST (Altschul et al., 1997; Schäffer et al., 2001) (maximum e-value = 1e^-4^; enable short queries and filtering of low complexity regions). Retrieved hits were collapsed using CD-HIT (Huang et al., 2010) with minimum sequence identity = 0.97 to remove redundant sequences. The resulting sequences were merged with those used for phylogenetic analysis in (Greer et al., 2015) using MAFFT (--add) (Katoh et al., 2017; Kuraku et al., 2013). Gaps and duplicate sequences were removed from the merged alignment. Only sequences corresponding to the taxa in Figure 2B were retained. The alignment was then used for phylogenetic tree construction using MrBayes in the CIPRES Science Gateway (Miller et al., 2010) with 5 × 10^6^ generations. Protein sequences used for MrBayes analysis are given in Table S3.

#### Preparation of nuclear extracts with DNA methyltransferase activity

Vegetative *Tetrahymena* cells were grown in SSP medium to log-phase (~3.5 × 10^5^ cells/mL) and collected by centrifugation at 2,300 × g for 5 min in an SLA-3000 rotor. The supernatant was discarded, and cells were resuspended in medium B (10mM Tris pH 6.75, 2mM MgCl_2_, 0.1M sucrose, 0.05% w/v spermidine trihydrochloride, 4% w/v gum Arabic, 0.63% w/v 1-octanol, and 1mM PMSF). Gum arabic (Sigma Aldrich) is prepared as a 20% w/v stock and centrifuged at 7,000 × g for 30 min to remove undissolved clumps. For each volume of cell culture, one-third volume of medium B was added to the *Tetrahymena* cell pellet. Cells were resuspended and homogenized in a chilled Waring Blender (Waring PBB212) at high speed for 40 sec. The resulting lysate was subsequently centrifuged at 2,750 × g for 5 min in an SLA-3000 rotor to pellet macronuclei. The nuclear pellet was washed twice with medium B, five times in MM medium (10mM Tris-HCl pH 7.8, 0.25M sucrose, 15mM MgCl_2_, 0.1% w/v spermidine trihydrochloride, 1mM DTT, 1mM PMSF). Macronuclei were pelleted between wash steps by centrifuging at 2,500 × g for 5 min in an SLA-3000 rotor. Finally, the total number of washed macronuclei was counted with a hemocytometer using a Zeiss ID03 microscope. Nuclear proteins were extracted by vigorously resuspending the pellet in MMsalt buffer (10mM Tris-HCl pH 7.8, 0.25M sucrose, 15mM MgCl_2_, 350mM NaCl, 0.1% w/v spermidine trihydrochloride, 1mM DTT, 1mM PMSF). 1mL MMsalt buffer was added per 2.33 × 10^8^ macronuclei. The viscous mixture was nutated for 45 min at 4°C, and then cleared at 175,000 × g for 30 min at 4°C in a SW 41 Ti rotor. Following this, the supernatant was dialyzed in a Slide-A-Lyzer 3.5K MWCO cassette (Thermo Fisher) overnight at 4°C against two changes of MMminus medium (10mM Tris-HCl pH 7.8, 15mM MgCl_2_, 1mM DTT, 0.5mM PMSF). The dialysate was then centrifuged at 7,197 × g for 1 hr at 4°C to remove precipitates, and dialyzed overnight in a Slide-A-Lyzer 3.5K MWCO cassette (Thermo Fisher) at 4°C against two changes of MN3 buffer (30mM Tris-HCl pH 7.8, 1mM EDTA, 15mM NaCl, 20% v/v glycerol, 1mM DTT, 0.5mM PMSF). The final dialysate was cleared by centrifugation at 7,197g for 1.5 hr at 4°C, flash frozen, and stored at −80°C. This nuclear extract was used for all subsequent biochemical fractionation and 6mA methylation assays.

#### Partial purification of MTA1 from nuclear extracts

*Tetrahymena* nuclear extracts were passed through a HiTrap Q HP column (GE Healthcare) and eluted using a linear gradient of 15mM to 650mM NaCl in 30mM Tris-HCl pH 7.8, 1mM EDTA, 20% v/v glycerol, 1mM DTT, 0.5 mM PMSF, over 30 column volumes. Each fraction was assayed for DNA methyltransferase activity using radiolabeled SAM as described in the next section. The DNA methyltransferase activity eluted in two peaks, at ~60mM and ~365mM NaCl, termed the “low salt sample” and “high salt sample”. Fractions corresponding to each peak were pooled and passed through a HiTrap Heparin HP column (GE Healthcare). Bound proteins were eluted using a linear gradient of 60 mM to 1M NaCl (for the low salt sample) or 350mM to 1M NaCl (for the high salt sample) over 30 column volumes. Fractions with DNA methyltransferase activity were respectively pooled and dialyzed into 10mM sodium phosphate pH 6.8, 100mM NaCl, 10% v/v glycerol, 0.3mM CaCl_2_, 0.5mM DTT (for the low salt sample); or 30mM Tris-HCl pH 7.8, 1mM EDTA, 200mM NaCl, 10% v/v glycerol, 1mM DTT, 0.2mM PMSF (for the high salt sample). The dialyzed low salt sample was passed through a Nuvia cPrime column (Bio-Rad) and eluted using a linear gradient of 100 mM to 1M NaCl in 50 mM sodium phosphate pH 6.8, 10% (v/v) glycerol, 0.5 mM DTT. Separately, the dialyzed high salt sample was fractionated using a Superdex 200 10/300 GL column (GE Healthcare) in 30mM Tris-HCl pH 7.8, 1mM EDTA, 200mM NaCl, 10% v/v glycerol, 1mM DTT. Fractions from the Nuvia cPrime and Superdex 200 columns were dialyzed into 30mM Tris-HCl pH 7.8, 1mM EDTA, 15mM NaCl, 20% v/v glycerol, 1mM DTT, 0.5mM PMSF and assayed for DNA methyltransferase activity. Those with qualitatively low, medium, and high activity were subjected to mass spectrometry to identify candidate methyltransferase proteins (Figure 2G).

#### DNA methyltransferase assay

To generate the DNA substrate for this assay, a 954bp fragment was amplified by PCR from *Tetrahymena thermophila* strain SB210 macronuclear SB210 genomic DNA using PCR primers metGATC_F2 and metGATC_R2 (Table S4). The resulting product was purified using Ampure XP beads (Beckman Coulter). This 954bp region of the genome contains a high level of 6mA *in vivo*. Thus, the underlying DNA sequence may be intrinsically amenable to methylation by *Tetrahymena* MTA1. Note that the amplified product is devoid of DNA methylation as unmodified dNTPs were used for PCR.

Radioactive methyltransferase assay: 2.18µg DNA substrate was mixed with 4-8µl nuclear extract and 0.64µM ^3^H-labeled S-adenosyl-L-methionine ([^3^H]SAM) in 33mM Tris-HCl pH 7.5, 6mM EDTA, 4.3mM BME, in a 15µl reaction volume. Samples were incubated overnight at 37°C, and subsequently spotted onto 1cm × 1cm squares of Hybond-XL membrane (GE Healthcare). Membranes were then washed thrice with 0.2M ammonium bicarbonate, once with distilled water, twice with 100% ethanol, and finally air-dried for 1 hr. Each membrane was immersed in 5mL Ultima Gold (PerkinElmer) and used for scintillation counting on a TriCarb 2910 TR (PerkinElmer). This assay was used in Figures 2D and 2E.

Non-radioactive methyltransferase assay: 5.5µg DNA substrate was mixed with 2µl nuclear extract and 0.2mM S-adenosyl-L-methionine (NEB) in 33mM Tris-HCl pH 7.5, 6mM EDTA, 4.3mM BME in a 15µl reaction volume and incubated at 37°C overnight. DNA was then purified using a MinElute purification kit (QIAGEN), denatured at 95°C for 10 min, and snap cooled in an ice water bath. Samples were spotted on a Hybond N+ membrane (GE Healthcare), air dried for 5 min and UV-cross-linked with 120,000 µJ / cm^2^ exposure using an Ultra-Lum UVC-515 Ultraviolet Multilinker. The cross-linked membrane was blocked in 5% milk in TBST (containing 0.1% v/v Tween) and incubated with 1:1,000 anti-N6-methyladenosine antibody (Synaptic Systems) at 4°C overnight. The membrane was then washed three times with TBST, incubated with 1:3,000 Goat anti-rabbit HRP antibody (Bio-Rad) at room temperature for 1 hr, washed another three times with 1x TBST, and developed using Amersham ECL Western Blotting Detection Kit (GE Healthcare). This dot blot assay was used to measure 6mA levels in Figures 2F, 3B, and 5C.

#### Quantitative mass spectrometry analysis of dA and 6mA

10.5µg *Oxytricha* or *Tetrahymena* macronuclear genomic DNA was first digested to nucleosides by mixing with 14µl DNA degradase plus enzyme (Zymo Research) in a 262.5µl reaction volume. Samples were incubated at 37°C overnight, then 70°C for 20 min to deactivate the enzyme.

The internal nucleoside standards ^15^N_5_-dA and D_3_-6mA were used to quantify endogenous dA and 6mA levels in ciliate DNA. ^15^N_5_-dA was purchased from Cambridge Isotope Laboratories, while D_3_-6mA was synthesized as described in the following section. Nucleoside samples were spiked with 1 ng/µl ^15^N_5_-dA and 200 pg/µl D_3_-6mA in an autosampler vial. Samples were loaded onto a 1mm × 100mm C18 column (Ace C18-AR, Mac-Mod) using a Shimadzu HPLC system and PAL auto-sampler (20µl / injection) at a flow rate of 70µl / min. The column was connected inline to an electrospray source couple to an LTQ-Orbitrap XL mass spectrometer (Thermo Fisher). Caffeine (2 pmol/µl in 50% Acetonitrile with 0.1% FA) was injected as a lock mass through a tee at the column outlet using a syringe pump at 0.5µl/min (Harvard PHD 2000). Chromatographic separation was achieved with a linear gradient from 10% to 99% B (A: 0.1% Formic Acid, B: 0.1% Formic Acid in Acetonitrile) in 5 min, followed by 5 min wash at 100% B and equilibration for 10 min with 1% B (total 20 min program). Electrospray ionization was achieved using a spray voltage of 4.50 kV aided by sheath gas (Nitrogen) flow rate of 18 (arbitrary units) and auxiliary gas (Nitrogen) flow rate of 2 (arbitrary units). Full scan MS data were acquired in the Orbitrap at a resolution of 60,000 in profile mode from the m/z range of 190-290. A parent mass list was utilized to acquire MS/MS spectra at a resolution of 7500 in the Orbitrap. LC-MS data were manually interpreted in Xcalibur’s Qual browser (Thermo, Version 2.1) to visualize nucleoside mass spectra and to generate extracted ion chromatograms by using the theoretical [M+H] within a range of ±2ppm. Peak areas were extracted in Skyline (Ver. 3.5.0.9319).

#### Synthesis of D_3_-6mA nucleoside

2’-Deoxyadenosine and CD_3_I were purchased from Sigma-Aldrich. Flash chromatography was performed on a Biotage Isolera using silica columns (Biotage SNAP Ultra, HP-Sphere 25µm). Semi-preparative RP-HPLC was performed on a Hewlett-Packard 1200 series instrument equipped with a Waters XBridge BEH C18 column (5μm, 10 × 250 mm) at a flow rate of 4mL/min, eluting using A (0.1% formic acid in H2O) and B (0.1% formic acid in 9:1 MeCN/H2O). ^1^H NMR spectra were recorded on a Bruker UltraShield Plus 500 MHz instrument. Data for ^1^H NMR are reported as follows: chemical shift (δ ppm), multiplicity (s = singlet, br = broad signal, d = doublet, dd = doublet of doublets) and coupling constant (Hz) where possible. ^13^C NMR spectra were recorded on a Bruker UltraShield Plus 500 MHz.

D_3_-6mA (2’Deoxy-6-[*D*3]-methyladenosine) were synthesized and purified according to (Schiffers et al., 2017). After an initial purification by flash column chromatography, the methylated compounds were further purified by semipreparative RP-HPLC (linear gradient of 0% to 20% B over 30 min) affording the desired compounds in 14% and 10% yields respectively after lyophilization.

##### 2’Deoxy-6-[*D*3]-methyladenosine

^1^H NMR (500 MHz, D_2_O) δ 7.98 (s, 1H), 7.77 (s, 1H), 6.17 (m, 1H), 4.54 (m, 1H), 4.10 (m, 1H), 3.79 (dd, *J* = 12.7, 3.2 Hz, 1H), 3.71 (dd, *J* = 12.7, 4.3 Hz, 1H), 2.60 (m, 1H), 2.44 (ddd, *J* = 14.0, 6.3, 3.3 Hz, 1H).

^13^C NMR (126 MHz, D_2_O) δ 154.0, 151.5, 146.1, 138.9, 118.4, 87.3, 84.3, 71.1, 61.6, 39.2, 26.4 ppm. (Peak at 26.4 ppm appears as a broad signal. C-D coupling is not resolved).

HR-MS (ESI+): m/z calculated for [C_11_H_13_D_3_N_5_O_3_]^+^ ([M+H]^+^): 269.1436, found 269.1421.

#### Mass spectrometry analysis of proteins in *Tetrahymena* nuclear extracts

Samples where topped up to 200µl with 50mM ammonium bicarbonate pH 8. TCEP was added to 5mM final concentration and left to incubate at 60°C for 10 min. 15mM chloroacetamide was then added and left to incubate in the dark at room temperature for 30 min. 1µg of Trypsin Gold (Promega) was added to each sample and incubated end-over-end at 37°C for 16 hr. An additional 0.25µg of Trypsin Gold was added and incubated end-over-end at 37°C for 3 hr. Samples were acidified by adding TFA to 0.2% final concentration, and desalted using SDB stage-tips (Rappsilber et al., 2007). Samples were dried completely in a speedvac and resuspended in 20µl of 0.1% formic acid pH 3. 5µl was injected per run using an Easy-nLC 1200 UPLC system. Samples were loaded directly onto a 45cm long 75µm inner diameter nano capillary column packed with 1.9um C18-AQ (Dr. Maisch, Germany) mated to metal emitter in-line with an Orbitrap Fusion Lumos (Thermo Scientific, USA). The mass spectrometer was operated in data dependent mode with the 120,000 resolution MS1 scan (AGC 4e5, Max IT 50ms, 400-1500 m/z) in the Orbitrap followed by up to 20 MS/MS scans with CID fragmentation in the ion trap. Dynamic exclusion list was invoked to exclude previously sequenced peptides for 60 sec if sequenced within the last 30 sec, and maximum cycle time of 3 sec was used. Peptides were isolated for fragmentation using the quadrupole (1.6 Da window). ns was utilized. Ion-trap was operated in Rapid mode with AGC 2e3, maximum IT of 300 msec and minimum of 5000 ions.

Raw files were searched using Byonic (Bern et al., 2012) and Sequest HT algorithms (Eng et al., 1994) within the Proteome Discoverer 2.1 suite (Thermo Scientific, USA). 10ppm MS1 and 0.4Da MS2 mass tolerances were specified. Caramidomethylation of cysteine was used as fixed modification, while oxidation of methionine, pyro-Glu from Gln and deamidation of asparagine were specified as dynamic modifications. Trypsin digestion with maximum of 2 missed cleavages were allowed. Files were searched against the *Tetrahymena themophila* macronuclear reference proteome (June 2014 build), supplemented with common contaminants (27,099 total entries).

Scaffold (version Scaffold_4.8.7, Proteome Software Inc., Portland, OR) was used to validate MS/MS based peptide and protein identifications. Peptide identifications were accepted if they could be established at greater than 93.0% probability. Peptide Probabilities from Sequest and Byonic were assigned by the Scaffold Local FDR algorithm. Protein identifications were accepted if they could be established at greater than 99.0% probability to achieve an FDR less than 1.0% and contained at least 3 identified peptides. Protein probabilities were assigned by the Protein Prophet algorithm (Nesvizhskii et al., 2003). Proteins that contained similar peptides and could not be differentiated based on MS/MS analysis alone were grouped to satisfy the principles of parsimony.

#### Generation of *mta1* mutant lines

A frameshift mutation in the MTA1 gene was created by inserting a small non-coding DNA segment immediately downstream of the MTA1 start codon (Figures 3A and S6). This non-coding DNA segment belongs to a class of genetic elements that are normally eliminated during the sexual cycle (Chen et al., 2014). When ssRNA homologous to such DNA segments is injected into *Oxytricha* cells undergoing sexual development, the DNA is erroneously retained (Khurana et al., 2018). This results in disruption of the MTA1 open reading frame. The ectopic DNA segment is propagated through subsequent cell divisions after completion of the sexual cycle.

ssRNA was generated by *in vitro* transcription using a Hi-Scribe T7 High Yield RNA Synthesis Kit (NEB). The DNA template for *in vitro* transcription consists of the ectopic DNA segment flanked by 100-200bp cognate MTA1 sequence. Following DNase treatment, ssRNA was acid-phenol:chloroform extracted and ethanol precipitated. After precipitation, ssRNA was resuspended in nuclease-free water (Ambion) to a final concentration of 1 to 3 mg/mL for injection.

#### ssRNA microinjections

*Oxytricha* cells were mated by mixing 3mL of each mating type, JRB310 and JRB510, along with 6mL of fresh Pringsheim media. At 10 to 12 hr post mixing, pairs were isolated and placed in Volvic water with 0.2% bovine serum albumin (Jackson ImmunoResearch Laboratories) according to previously published methods (Fang et al., 2012). ssRNA constructs were injected into the macronuclei of the pair cells as previously described with DNA constructs (Nowacki et al., 2008). After injection, cells were pooled in Volvic water. At 60 to 72 hr post mixing, the pooled cells were singled out to grow clonal injected cell lines. As clonal population size grew, lines were transferred to 10 cm petri dishes and grown in Pringsheim media.

#### Quantification and statistical analysis

All statistical tests were performed in Python (v2.7.10) or R (v3.2.5), and described in the respective Figure and Table legends.

#### Data availability

*Oxytricha* SMRT-seq data are deposited in SRA under the accession numbers SRX2335608 and SRX2335607. *Tetrahymena* SMRT-seq and all *Oxytricha* Illumina data are deposited in NCBI GEO under accession number GSE94421 (private link: *https://www.ncbi.nlm.nih.gov/geo/query/acc.cgi?token=qbcneqcolbkrrkp&acc=GSE94421*).

